# The centrosome is a selective phase that nucleates microtubules by concentrating tubulin

**DOI:** 10.1101/093054

**Authors:** Jeffrey B. Woodruff, Beatriz Ferreira Gomes, Per O. Widlund, Julia Mahamid, Anthony A. Hyman

## Abstract

Centrosomes are non-membrane-bound compartments that nucleate microtubule arrays. They consist of nanometer-scale centrioles surrounded by a micron-scale, dynamic assembly of protein called the pericentriolar material (PCM). To study how PCM forms a spherical compartment that nucleates microtubules, we reconstituted PCM-dependent microtubule nucleation *in vitro* using recombinant *C.elegans* proteins. We found that macromolecular crowding drives phase separation of the key PCM scaffold protein SPD-5 into spherical droplets that morphologically and dynamically resemble *in vivo* PCM. These SPD-5 droplets recruited the microtubule polymerase ZYG-9 (XMAP215 homologue) and the microtubule-stabilizing protein TPXL-1 (TPX2 homologue). Together, these three proteins concentrated tubulin ~4- fold over background, which was sufficient to reconstitute nucleation of microtubule asters *in vitro.* Our results suggest that *in vivo* PCM is a selective phase that organizes microtubule arrays through localized concentration of tubulin by microtubule effector proteins.

**One Sentence Summary:** Phase separation of *C. elegans* centrosome proteins drive the formation of micron-sized microtubule organizing centers.

## Introduction

Centrosomes are major microtubule-organizing centers (MTOCs) that play important roles in mitotic spindle assembly, asymmetric cell division, and polarity. A centrosome consists of centrioles that organize an amorphous assembly of protein called the pericentriolar material (PCM) that serves to nucleate microtubules. During interphase, the mother centriole organizes a thin, patterned layer (~200 nm) of PCM (Fu and Glover, 2012; Lawo et al., 2012; Mennella et al., 2012; Sonnen et al., 2012). As cells prepare for mitosis, a more expansive and less ordered PCM layer accumulates that can reach several microns in diameter. Decades of genetics, cell biology, and biochemistry have identified key proteins required for PCM assembly (for reviews, see (Conduit et al., 2015; Woodruff et al., 2014)). It is now understood that coiled-coil proteins, such as Cdk5Rap2 (vertebrates), Centrosomin *(Drosophila)* and SPD-5 (*C. elegans)*, self-assemble to form the underlying PCM scaffold onto which all other PCM proteins are loaded (Conduit et al., 2014a; Fong et al., 2008; Hamill et al., 2002; Megraw et al., 1999; Woodruff et al., 2015). The formation of these expansive scaffolds during mitotic entry requires Polo-like kinase, and SPD-2/Cep192 (Conduit et al., 2014a; 2014b; Dix and Raff, 2007; Giansanti et al., 2008; Kemp et al., 2004; Pelletier et al., 2004; Sumara et al., 2004; Sunkel and Glover, 1988; Woodruff et al., 2015; Zhu et al., 2008).

How coiled-coil PCM proteins form a scaffold that organizes microtubule asters is not understood. In fact, the minimal requirements to organize microtubule arrays are unknown in any system. The classical viewpoint proposes that PCM serves as a binding platform for gamma tubulin–containing complexes that template microtubule nucleation (Moritz et al., 1995; Schnackenberg et al., 1998; Zheng et al., 1995). Kollman *et al.* (2015) demonstrated that yeast gamma tubulin small complexes efficiently nucleate microtubules *in vitro*, supporting this idea. However, knock down of gamma tubulin *in vivo* reduces, but does not fully eliminate, PCM-based microtubule nucleation (Hannak et al., 2002; Sampaio et al., 2001; Strome et al., 2001; Sunkel et al., 1995), indicating that additional mechanisms exist to nucleate microtubules. An alternative nucleation pathway might involve ch-TOG family proteins (XMAP215 in frogs and ZYG-9 in *C. elegans*) and/or TPX2 family proteins (TPXL-1 in *C. elegans)*, as they localize to PCM and regulate microtubule dynamics both in living cells and in reconstituted systems (Brouhard et al., 2008; Brunet et al., 2004; Gergely et al., 2003; Matthews et al., 1998; Özlü et al., 2005; Roostalu et al., 2015; Wieczorek et al., 2015). In addition to recruiting the right proteins, PCM must also possess certain material properties to function properly as a MTOC. PCM must be porous enough to permit entry and diffusion of microtubule-associated proteins and tubulin dimers, while still retaining these molecules. Furthermore, PCM must be flexible enough to allow PCM expansion, while being strong enough to resist microtubule-dependent pulling forces. We still do not understand how PCM forms a dynamic scaffold that can concentrate microtubule-associated proteins and tubulin to form a robust microtubule aster, in part due to a lack of a reconstituted system to study such questions.

The *C. elegans* embryo is an excellent system to study PCM-based microtubule nucleation. Genetic experiments have identified a core set of proteins that are essential for the PCM assembly: SPD-2/CEP192, Polo–like kinase 1 (PLK-1), and the coiled-coil protein SPD-5 (Decker et al., 2011; Hamill et al., 2002; Kemp et al., 2004; Pelletier et al., 2004). The simplicity of the *C. elegans* PCM suggests that it may be possible to reconstitute PCM-dependent microtubule nucleation *in vitro.* Indeed, purified SPD-5 self-assembles into micron-sized scaffolds capable of specifically recruiting downstream PCM proteins *in vitro* (Woodruff et al., 2015). SPD-2 and PLK-1 regulate the formation of these scaffolds, consistent with their roles in driving PCM expansion *in vivo* (Woodruff et al., 2015). However, these in *vitro* SPD-5 scaffolds are extended assemblies with no regular shape, unlike *in vivo* centrosomes, which are spherical. More importantly, they are unable to nucleate microtubules (Woodruff et al., 2015).

Here, we use biochemical reconstitution to identify a minimal system that achieves proper centrosome morphology and microtubule organization. We present evidence that cytoplasmic crowding guides SPD-5 assembly into a concentrated droplet state that resembles *C. elegans* PCM *in vivo.* These droplets efficiently recruit microtubule-nucleating proteins and tubulin and, as a result, are sufficient for the self-organization of micron-scale MTOCs.

## RESULTS

### SPD-5 molecules self-organize into micron-sized, amorphous droplets in a crowded environment *in vitro*

In non-physiological conditions containing high detergent, high salt, and cold temperatures, SPD-5 protein is monomeric. When diluted into buffer containing low salt and no detergent and incubated at room temperature, SPD-5 assembles into micron-scale, supramolecular networks with no regular shape or higher-order structure (Woodruff et al., 2015). Although SPD-5 networks are subject to physiological regulation and serve as scaffolds for PCM proteins (Woodruff et al., 2015), they are dispersed structures that do not match the dense, spherical morphology of *in vivo* PCM (Figure 1A). Furthermore, they do not nucleate microtubules (data not shown).

**Figure 1.**
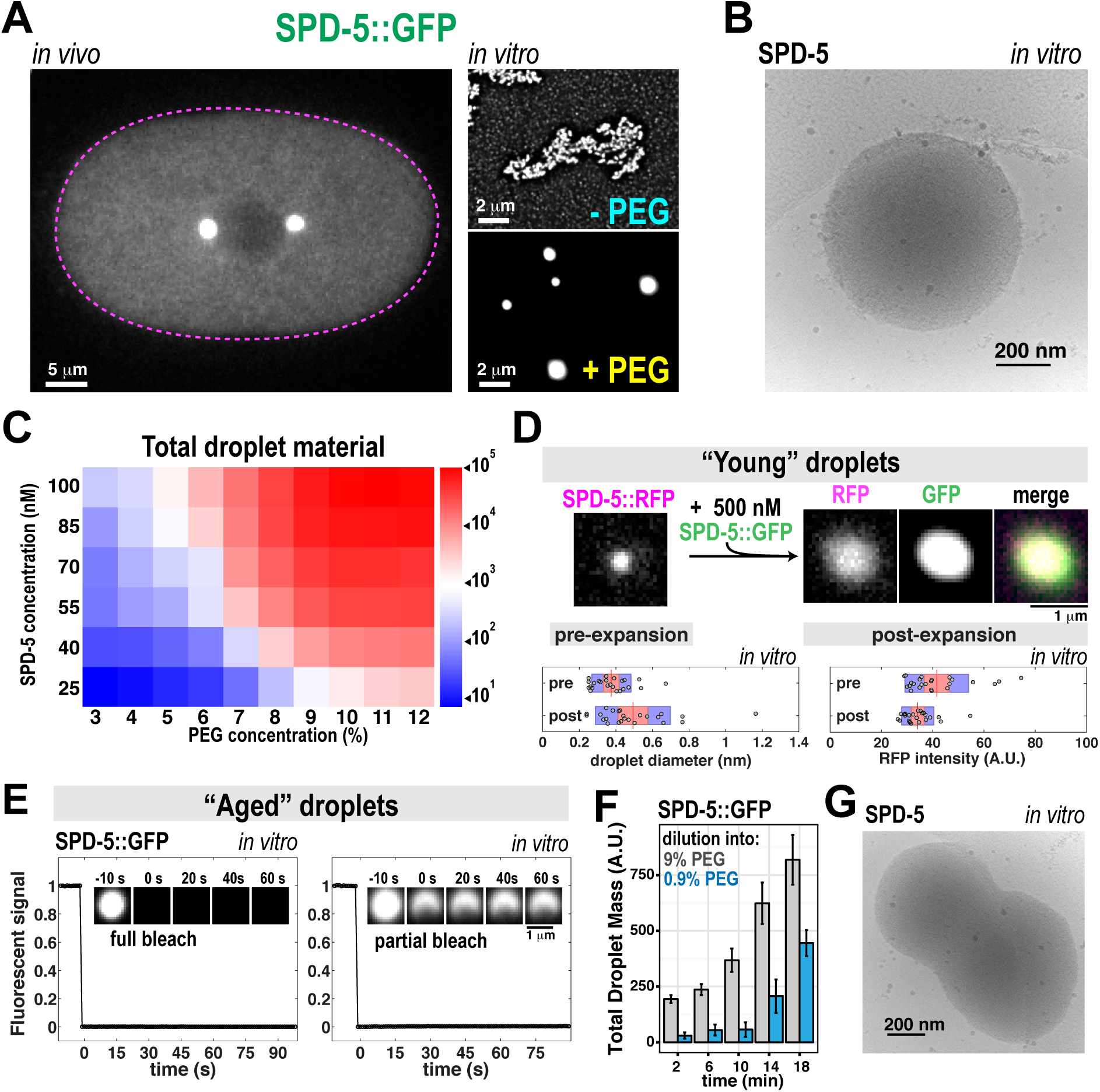
Purified SPD-5 forms micron-scale droplets in the presence of a macromolecular crowding agent *in vitro.* A. GFP-labeled SPD-5 assembles into spherical pericentriolar material in a *C. elegans* embryo (left). Cell outline is in magenta. Purified SPD-5::GFP incubated without polyethylene glycol (-PEG) assembles into networks, whereas SPD-5::GFP forms micron-scale droplets in the presence of >4% (w/v) polyethylene glycol (+PEG).
B. Cryo electron microscopy image of a SPD-5 droplet in 9% PEG.
C. High-throughput analysis of droplet formation in 60 different conditions. The color indicates the average amount of droplet material per image after ten minutes of incubation at 23°C. All conditions were performed in quadruplicate with 6 images taken per replicate (1,440 images total).
D. Small SPD-5 droplet seeds were formed by incubating 25 nM SPD-5::TagRFP in 9% PEG for 30 s ("young” droplets), then expanded by adding 500 nM SPD-5::GFP. Images on the right were taken 4.5 min after expansion. The RFP signal grew in area but decreased in intensity after expansion (p =0.03 and p =0.01, respectively; Wilcoxon rank sum test; n = 24 droplets per condition), indicating internal rearrangement. Plots show means (red lines), 95% confidence intervals (red shaded areas), and S.D. (blue shaded areas).
E. 500 nM SPD-5::GFP droplets were imaged 15 min after formation ("aged” droplets). Fluorescent signal did not recover after photobleaching the entire droplet (n = 15) or part of the droplet (n = 17).
F. Droplet dilution assay. 500 nM SPD-5::GFP droplets were formed in 9% PEG for various lengths of time, then diluted 10-fold into buffer containing 9% or 0% PEG (final concentration = 0.9% PEG). Total droplet mass per field of view was calculated (mean ± 95% confidence intervals; n = 11 experiments per time point).
G. Cryo electron microscopy image of coalescing SPD-5 droplets frozen 2 min after droplet formation. See also Figure S1.

To identify the minimal factors that make a centrosome competent to organize microtubules, we first considered the effect of the cytoplasm on PCM assembly. The interior of a cell is extremely dense and viscous, containing ~80-400 mg/ml macromolecules (proteins, nucleic acids, polysaccharides, etc.) (van den Berg et al., 1999; Zimmerman and Trach, 1991). Typical “physiological” buffers match the pH and salt concentrations found *in vivo*, yet these buffers do not have a crowding effect. We generated a minimal crowded environment by adding polyethylene glycol (PEG) to assembly reactions containing physiological concentrations of SPD-5 (500 nM; Figure S1A)(Saha et al., 2016). In the presence of >4% PEG (molecular weight (MW): 3,350 Da), SPD-5::GFP formed micron-sized, round assemblies similar to the size and shape of SPD-5-labeled PCM *in vivo* (Figure 1A). Cryo-electron microscopy (EM) revealed that these SPD-5 assemblies were spherical and amorphous (Figure 1B). Using high-throughput automated imaging and segmentation to test 60 different combinations of PEG and SPD-5 concentrations, we constructed a phase diagram which showed that the mass and number of spherical SPD-5 assemblies increased with either PEG or SPD-5 concentration (Figure 1C and S1B). Other proteins, such as PLK-1::GFP, TPXL-1::GFP, EB1-GFP, and GFP, did not form similar structures under similar conditions (Figure S1C; all proteins are shown in Figure S1D).

In *C. elegans* embryos, PCM rapidly expands in preparation for mitosis, then stops prior to the metaphase-anaphase transition (Decker et al., 2011). During PCM growth SPD-5 incorporates throughout the PCM, indicating that PCM expands isotropically (Laos et al., 2015). *In vitro* SPD-5 assemblies also grew by isotropic incorporation and expansion. When “young” (incubated for < 2 min) SPD-5::TagRFP seeds (25 nM) were diluted into a solution containing unassembled SPD-5::GFP (500 nM), SPD-5::GFP incorporated into the seeds. The seeds subsequently expanded, on average, ~25% in diameter (p = 0.03, n = 24; Wilcoxon rank sum test), and the RFP signal became concomitantly ~20% dimmer (p = 0.01), indicating that SPD-5 originally in the seeds redistributed within the growing assemblies (Figure 1D and S1E). Once PCM expansion stops during mitosis *in vivo*, SPD-5::GFP signal does not recover after full-bleach nor redistribute after half-bleach of the PCM (Laos et al., 2015). Similarly, FRAP analysis showed that “aged” (incubated for >10 min) SPD-5::GFP spherical assemblies also did not recover after full or partial bleaching, indicating that neither monomer incorporation nor long-range internal rearrangement occurs in this state (Figure 1E).

To further characterize the dynamic properties of *in vitro* SPD-5 spherical assemblies, we analyzed their size, number, reversibility, and ability to fuse. SPD-5 spherical assemblies grew over time even as their total number decreased (Figure S1F). Such behavior is known as Ostwald ripening, implying that smaller assemblies are unstable and disappear either through dissolution or fusion with other assemblies. Indeed, 10-fold dilution of young SPD-5 assemblies (500 nM, 2 min old) into PEG-free buffer (final concentration = 0.9% PEG) triggered their instantaneous dissolution. However, aged SPD-5 assemblies (18 min old) were 3.5-fold more resistant to dissolution after dilution (Figure 1E). Furthermore, we observed fusion of young SPD-5 spherical assemblies using cryo EM (Figure 1G; we could not assess fusion using light microscopy due to the small size of young assemblies). Aged SPD-5 assemblies merely clumped together (Figure S1G). These data suggest that the PCM scaffold is an evolving material that becomes less dynamic with time.

The fact that these *in vitro* SPD-5 assemblies are amorphous, spherical, initially labile, and coarsen, but become less dynamic with time suggests that they are viscous liquids that rapidly harden. This hardened structure could either be a gel or a glass (see Discussion). The phenomenon whereby a liquid-like state hardens into a more viscous material has been called “ageing” or “maturation”, and has been reported for other protein droplet systems as a pathological process (Lin et al., 2015; Patel et al., 2015). Our data suggest that ageing may serve a physiological role in the case of PCM. Based on these properties, we therefore call these dense, spherical SPD-5 assemblies “droplets” for the rest of the paper.

### Depletion attraction forces drive the formation of SPD-5 droplets

Purified SPD-5::GFP formed micron-scale droplets in the presence of other long-chain, inert polymers, such as Ficoll and Dextran, or in a highly concentrated solution of non-centrosomal protein, such as lysozyme (1 mM)(Figure S2A). Surprisingly, we found that addition of glycerol, another typical crowding agent, did not induce SPD-5 droplet formation (Figure S2A). Why would long-chain polymers (MW: >3,350 Da) and protein (MW: 14,300 Da) influence SPD-5 assembly, but not a small molecule like glycerol (MW: 92 Da)? These molecules are predicted to be inert, suggesting that the only difference between them is molecular mass. If the effect is truly due to size of the crowding agent, and not some unrecognized chemical effect, then changing the chain length of PEG should influence droplet formation. Thus, we incubated 500 nM SPD-5::TagRFP in solutions containing PEG molecules of varying average molecular mass (0.3-35 kDa). PEG concentration was kept constant at 9% (w/v) to generate similar crowding conditions. When incubated with 0.3-0.6 kDa PEG, SPD-5::TagRFP did not form droplets, but rather formed networks after 30 min, similar to when no crowding agent was used (Figure 2; Figure S2B). When larger PEGs (>1 kDa) were used as crowding agents, SPD-5::TagRFP assembled into droplets. Droplet formation efficiency peaked with 6 kDa PEG and dropped off at larger or smaller PEG sizes (Figure 2). These results indicate that SPD-5 droplet formation is sensitive to the size of the crowding agent. Because the effect has a peak and tails off for smaller and larger PEG sizes, it is likely that a depletion attraction mechanism drives SPD-5 assembly in the context of a crowded environment (see Discussion). For the remainder of our experiments, we used 3.35 kDa PEG for consistency.

**Figure 2.**
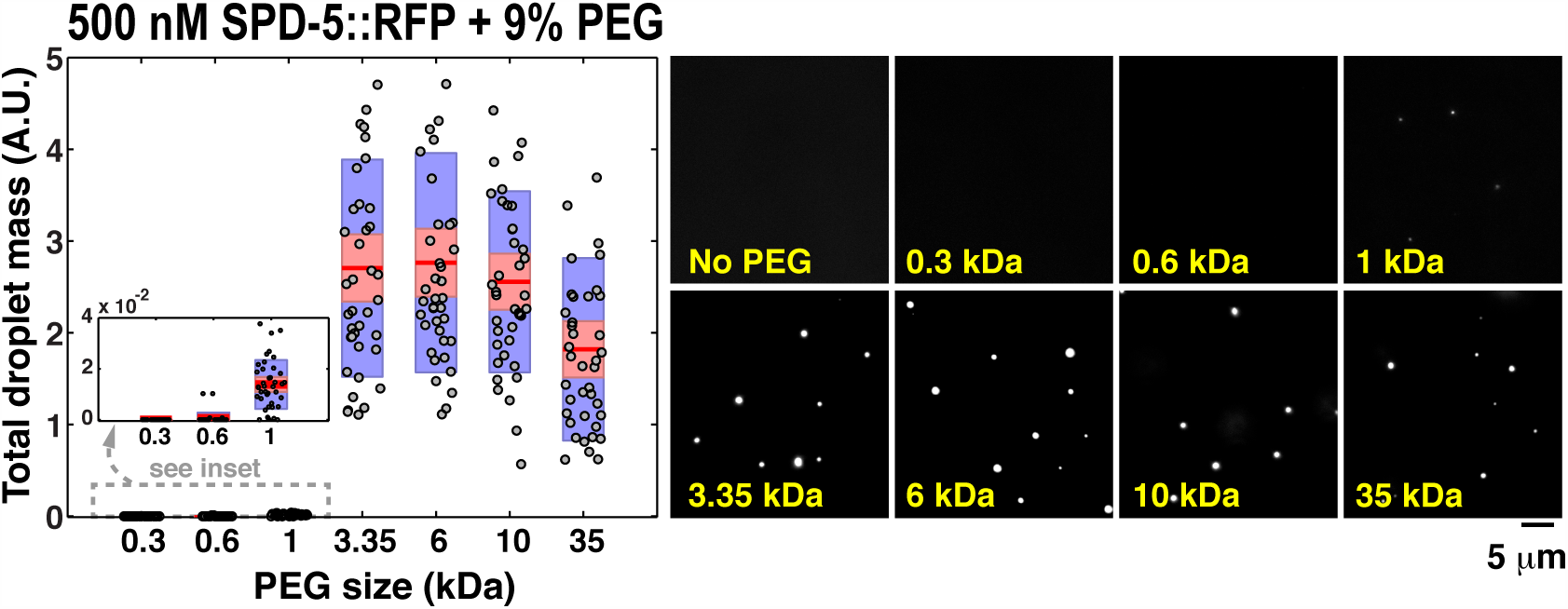
SPD-5 droplet formation is sensitive to the size of the crowding agent. 500 nM SPD-5::TagRFP was incubated in 9% solutions of PEG ranging in average polymer molecular weight (0.3-35 kDa) for 10 min, then analyzed by microscopy. Total droplet mass per image was calculated. Plot shows means (red lines), 95% confidence intervals (red shaded areas), and S.D. (blue shaded areas; n = 40 images per condition). See also Figure S2.

### PLK-1 and SPD-2 coordinately regulate SPD-5 phase separation

*In vivo*, PCM assembles around centrioles and is regulated by two PCM-localized proteins, the Polo-like Kinase PLK-1 and the Cep192 homologue SPD-2. Depletion or inactivation of these proteins prevents centrosomal accumulation of all known PCM proteins, including SPD-5 (Decker et al., 2011; Kemp et al., 2004; Pelletier et al., 2004; Woodruff et al., 2015). To study regulation, we used the phase diagram in Figure 1C to identify conditions in which spontaneous SPD-5 droplet formation does not occur (3% PEG and 100 nM SPD-5; Figure 3A). In this condition, addition of 100 nM PLK-1 or SPD-2 promoted SPD-5 droplet formation *in vitro* (Figure 3A,B).

**Figure 3.**
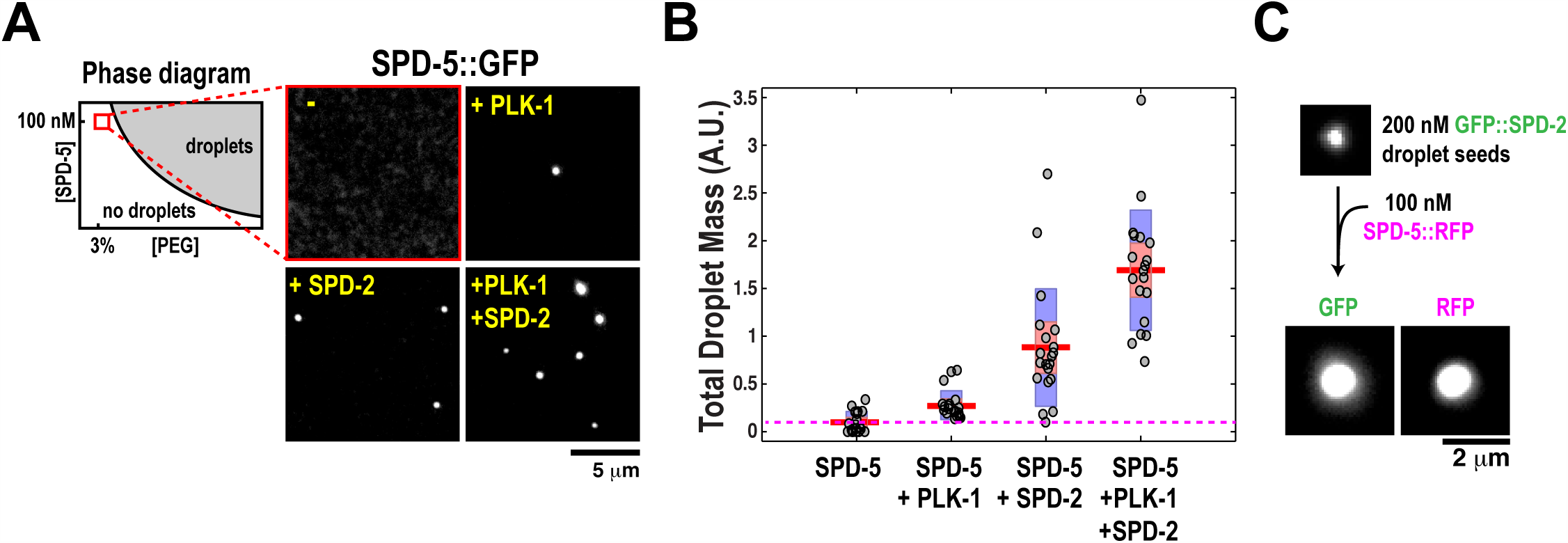
SPD-2 and PLK-1 regulate SPD-5 phase separation *in vitro.* A. 100 nM SPD-5::GFP was incubated a solution containing 3% PEG + 0.2 mM ATP and analyzed after 10 min at 23°C. Droplets do not form unless 100 nM SPD-2 or 100 nM PLK-1 is added. Phase diagram is a schematic of Figure 1C.
B. Quantification of (A). Total droplet mass per image was calculated. Plot shows means (red bars), 95% confidence intervals (red shaded areas), and S.D. (blue shaded areas; n = 20 images per condition).
C. 200 nM GFP::SPD-2 forms compact seeds when incubated 5% PEG. These seeds recruit SPD-5::TagRFP and grow into droplets (n > 50 seeds). See also Figure S3.

SPD-2 localizes to centrioles, is the most upstream component of the PCM recruitment hierarchy, and has been speculated to nucleate PCM (Kemp et al., 2004; Pelletier et al., 2004). We wondered if locally concentrated SPD-2 could act as a scaffold to trigger PCM nucleation. To mimic the localized nucleation of SPD-5 seen *in vivo*, we performed a “seeded assembly” experiment. We formed bright, diffraction-limited SPD-2 seeds by incubating 200 nM GFP::SPD-2 in 5% PEG (Figure S3A). When added to a solution of monomeric SPD-5::TagRFP (100 nM), these SPD-2 seeds triggered the formation of SPD-5::TagRFP droplets (Figure 3C). Using this seeding method, we could generate SPD-5 droplets PEG concentrations as low as 1.5% (Figure S3B). These results suggest that concentration of active SPD-2 around centrioles is a key step in nucleating formation of the SPD-5 scaffold *in vivo*.

### SPD-5 droplets are three-dimensional scaffolds that concentrate PCM client proteins

In embryos, genetic evidence has suggested that SPD-5 serves as a scaffold that recruits numerous downstream effector proteins such as PLK-1, SPD-2, TPXL-1, and ZYG-9, referred to hereafter as “PCM clients” (Hachet et al., 2007; Hamill et al., 2002; Kemp et al., 2004; Özlü et al., 2005; Pelletier et al., 2004). To compare the behavior of SPD-5 droplets with PCM, we measured partition coefficients (*PCs*), which we define as the ratio of client concentrations in the PCM vs. the cytoplasm (*in vivo*), or in droplets vs. the bulk phase (*in vitro*). *In vivo*, ZYG-9::mCherry (*PC* = 23 ± 9; mean ± S.D.; n = 33 centrosomes) was the most strongly recruited, followed by TPXL-1::GFP (*PC* = 18 ± 7), SPD-2::GFP (*PC* = 17 ± 6), and then PLK-1::GFP (*PC* = 2.8 ± 0.6)(Figure S4A). *In vitro*, partition coefficients were on average higher, but followed a similar trend. SPD-5 droplets recruited ZYG-9::GFP (*PC* = 101 ± 37; mean ± S.D.; n = 195 droplets) most efficiently, followed by GFP::SPD-2 (*PC* = 67 ± 41), TPXL-1::GFP (*PC* = 59 ± 33), and PLK-1::GFP (*PC* = 19.8 ± 7)(Figure 4A). *In vitro* droplets concentrated EB1-GFP weakly (*PC* = 3.2 ± 1) and did not concentrate GFP at all (*PC* = 1 ± 0.1). These data suggest that the SPD-5 scaffold *per se* determines the selectivity of the PCM for its clients. Moving to a complex system where scaffold binding capacity is more limited may be important for establishing proper *PCs in vitro*.

We also compared the mobility of the client proteins within *in vitro* SPD-5 droplets and *in vivo* PCM. We chose to analyze centrosomes in the P_2_ and EMS cells of 4-cell embryos because at this stage centrosome size and intensity are stable for several minutes after metaphase onset (Figure 4B; (Decker et al., 2011)). Consistent with previous analysis (Laos et al., 2015), we found that fluorescence recovery of GFP::SPD-5 was marginal after the entire centrosome was bleached (~14% recovery after 100s; Figure 4C). PLK-1::GFP signal recovered quickly (t ½ = 4.6s), whereas TPXL-1::GFP (t ½ = 22.8s), SPD-2::GFP (t ½ = 53.6s), and ZYG-9::mCherry (t ½ = 66.4s) signals recovered more slowly (Figure 4D). After photobleaching part of the PCM, TPXL-1::GFP, SPD-2::GFP, and ZYG-9::mCherry signals redistributed in a directional pattern from the inside-out, indicating internal rearrangement (Figure S4B-E). We could not perform the same analysis of PLK-1::GFP *in vivo* for technical reasons (see materials and methods). Thus, PCM clients exhibit different degrees of mobility within the PCM.

**Figure 4.**
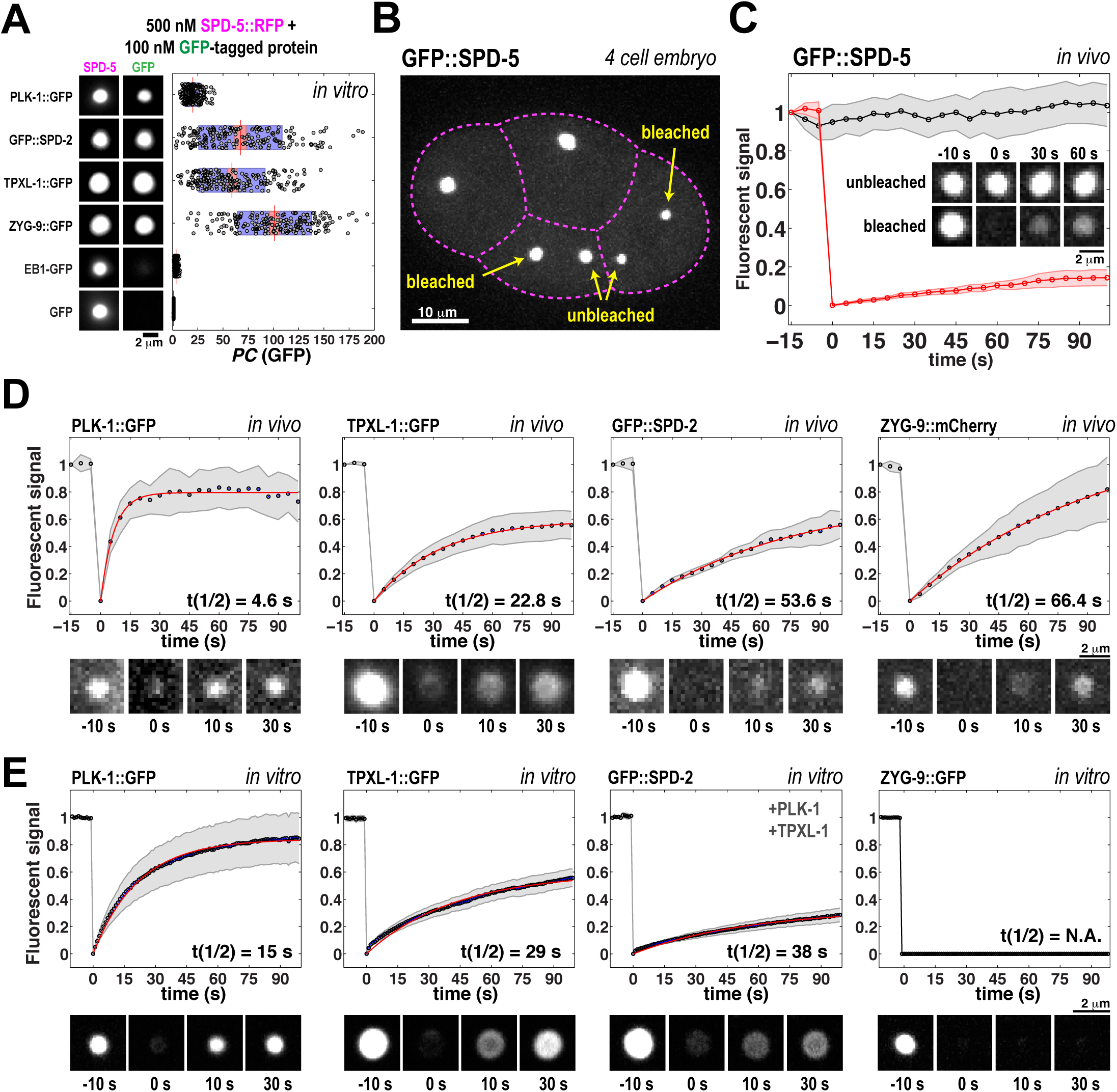
FRAP analysis of PCM client proteins *in vivo* and *in vitro*. A. 500 nM SPD-5::TagRFP was incubated for 1 min at 23°C in a 9% PEG solution, then 100 nM GFP-labeled proteins were added. Partition coefficients *(PC)* for the GFP-tagged proteins were analyzed after 10 min. Plots show means (red lines), 95% confidence intervals (red shaded areas), and S.D. (blue shaded areas; n = 195 droplets per condition).
B. Confocal microscope image of a 4-cell *C. elegans* embryo expressing GFP::SPD-5. For experiments in (C-D), the indicated centrosomes were photobleached (arrows).
C. Comparison of bleached (red line; n = 12) and unbleached (black line; n = 10) GFP::SPD-5-labeled centrosomes *in vivo.* Data are mean (dots) and 95% confidence intervals (shaded areas).
D. Fluorescence intensity recovery after photobleaching of *in vivo* centrosomes labeled with PLK-1::GFP (n= 10), TPXL-1::GFP (n = 12), GFP::SPD-2 (n = 10), and ZYG-9::mCherry (n = 12). Data are mean (dots) and 95% confidence intervals (shaded areas). The data were fit with a single exponential function to obtain half-life of recovery (t1/2, red line).
E. Fluorescence intensity recovery curves for *in vitro* SPD-5 droplets (500 nM) labeled with PLK-1::GFP (100 nM; n= 18), TPXL-1::GFP (1 μM; n = 19), GFP::SPD-2 (500 nM; n = 25), and ZYG-9::GFP (500 nM; n = 11). 250 nM of unlabeled PLK-1 and TPXL-1 were included in the GFP::SPD-2 experiment. See also Figure S4.

FRAP analysis indicated that PCM client proteins in SPD-5 droplets recovered with kinetics similar to that observed *in vivo* (Figure 4E). After photo-bleaching the entire droplet, the fluorescence intensity of PLK-1::GFP (t ½ = 15 s) recovered the fastest, followed by TPXL-1::GFP (t ½ = 29s), and GFP::SPD-2 (t ½ = 38s). Thus, these proteins freely exchange between the droplet and bulk environments. Partial photobleaching experiments indicated that these proteins also rearranged within SPD-5 droplets (Figure S4G-I). It must be noted that GFP::SPD-2 mobility required the presence of unlabeled PLK-1 and TPXL-1 (Figure 4E and S4F). So far, we have not observed ZYG-9 recovery or internal rearrangement within SPD-5 droplets under any circumstances (Figure 4E and S4J). PCM proteins, such as RSA-1, RSA-2 and TAC-1, which are missing from our reconstituted system, may be needed to further tune ZYG-9 properties. Nevertheless, the combination of *in vivo* and *in vitro* data suggest that PCM is a porous but selective phase composed of co-existing scaffold (low-turnover) and client (higher turnover) components.

### SPD-5/ZYG-9/TPXL-1 droplets concentrate tubulin *in vitro*

We next investigated whether SPD-5 droplets can also concentrate tubulin. We incubated 500 nM SPD-5::TagRFP droplets with 2 mM GTP and 200 μM alexa488-labeled tubulin for 10 min at 23°C. Under these conditions, SPD-5::TagRFP droplets did not concentrate tubulin, and MTs did not form in the bulk solution (Figure 5A). Addition of TPXL-1 or ZYG-9 made SPD-5 droplets competent to accumulate tubulin (*PC*_tubulin_ = 1.1 ± 0.1 (SPD-5 alone); 2 ± 0.5 (SPD-5 + TPXL-1); 3.1 ± 1 (SPD-5 + ZYG-9); 4 ± 2 (SPD-5 + TPXL-1 + ZYG-9); mean ± S.D.; n = 92 droplets). In SPD-5 droplets that contained ZYG-9, we often noticed bright tubulin puncta, which could represent subdomains within the larger droplet (Figure 5A). After partial photobleaching of SPD-5/TPXL-1/ZYG-9 droplets, alexa488-tubulin signal recovered in a directional pattern from the inside out (Figure 5B), indicating internal rearrangement.

**Figure 5.**
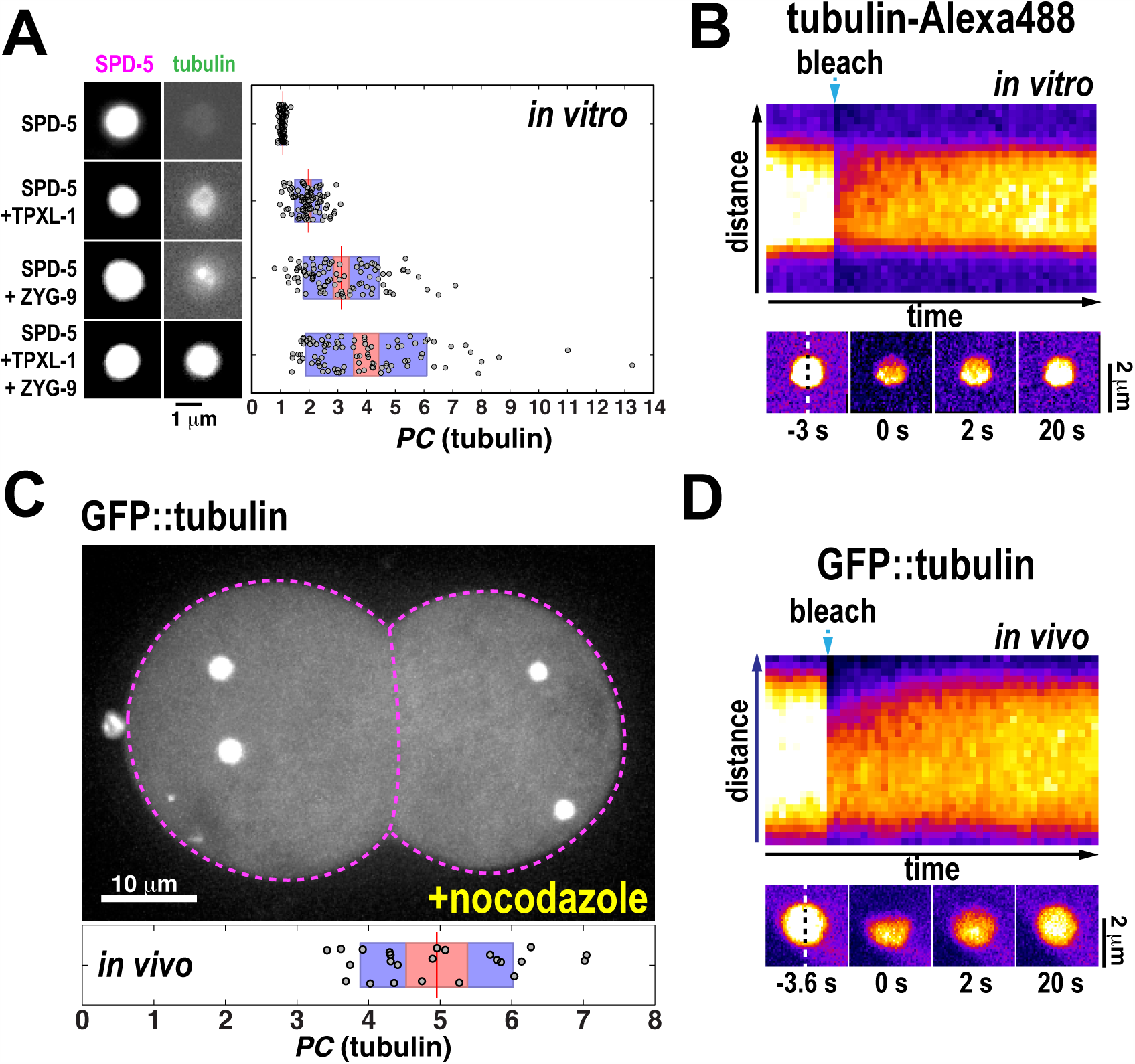
Pericentriolar material concentrates tubulin *in vitro and in vivo*. A. 500 nM SPD-5::TagRFP droplets were incubated with 200 nM α/β tubulin (alexa488-labeled), 2 mM GTP, 100 nM TPXL-1, and 50 nM ZYG-9. After 10 min, partition coefficients were measured for tubulin (PC(tubulin); n = 92 droplets). Plot shows means (red lines), 95% confidence intervals (red shaded areas), and S.D. (blue shaded areas).
B. Fluorescence recovery of alexa488-tubulin signal after partial bleaching of a SPD-5 droplet. The kymograph is along dotted line in the bottom image sequence. Images are pseudo-colored (white/yellow = high intensity; purple/black= low intensity). Note that signal recovers directionally starting from the unbleached region.
C. Permeable embryos expressing GFP-labeled beta tubulin (GFP::tubulin) were treated with nocodazole to depolymerize microtubules. Cell outline indicated by dotted magenta line. Ratio of centrosome to cytoplasmic fluorescence is shown (*PC*(tubulin)). Plot shows means (red bars), 95% confidence intervals (red shaded areas), and S.D. (blue shaded areas; n = 24 centrosomes).
D. Fluorescence recovery of GFP::tubulin signal in a nocodazole-treated embryo after partial bleaching of a centrosome. The kymograph is along the dotted line in the bottom image sequence.

We then examined the concentration of tubulin into PCM *in vivo.* To eliminate the contributions of microtubule polymerization to signal intensity, we treated permeable embryos with 20 μg/ml nocodazole (Carvalho et al., 2011). Microtubules depolymerized upon nocodazole treatment, but centrosomes still concentrated GFP::tubulin ~5-fold compared to the cytoplasm (*PC*_tubulin_ = 4.95 ± 1; n = 24 centrosomes; Figure 5C), similar to the partition coefficient of tubulin the SPD-5/TPXL-1/ZYG-9 droplets (*PC*_tubulin_ = 4; Figure 5A). Partial photobleaching of centrosomes revealed internal rearrangement of GFP::tubulin signal (Figure 5D), similar to *in vitro* droplets. Taken together, the *in vivo* and in *vitro* experiments suggest that the PCM acts a selective phase that, by concentrating the microtubule-associated proteins ZYG-9 and TPXL-1, can concentrate tubulin 4-5-fold.

### SPD-5/ZYG-9/TPXL-1 droplets generate microtubule asters

Our experiments so far show that SPD-5 droplets act as a selective phase to concentrate microtubule-associated proteins, which in turn concentrate tubulin. To test if SPD-5 droplets could organize a radial microtubule array, we increased the tubulin concentration to 2.5 μM in the presence of TPXL-1 and ZYG-9. After 6 min, MTs emerged from SPD-5 droplets that contained ZYG-9 (Figure 6A,B). Addition of both TPXL-1 and ZYG-9 dramatically enhanced MT nucleation from SPD-5 droplets, creating robust MT asters (Figure 6A-C). We never observed MTs emanating from SPD-5 or SPD-5/TPXL-1 droplets (Figure 6A,B). Thus, TPXL-1 and ZYG-9 synergistically promote MT nucleation. We conclude that SPD-5/TPXL-1/ZYG-9 droplets are sufficient to nucleate MTs and arrange them in a radial array (Figure 6D). This activity depended on the ability of ZYG-9 to bind tubulin dimers. Mutation of the contact sites between tubulin and the three TOG domains in ZYG-9 (ZYG-9^AA^)(Widlund et al., 2011) dramatically reduced tubulin recruitment to SPD-5 droplets (*PC*_tubulin_ = 1.2 ± 0.1; n = 92 droplets) and aster formation (Figure 6E-G). As an additional control, we tested aster formation when SPD-5 was not present. Without SPD-5, ZYG-9::mCherry formed aggregates that recruited TPXL-1 (Figure S5A,B). These particles nucleated single MTs but did not organize MT arrays (Figure S5C). These data suggest that ZYG-9 concentration *per se* is sufficient for MT nucleation, but the formation of robust asters requires concentration of both ZYG-9 and TPXL-1 by SPD-5 droplets.

**Figure 6.**
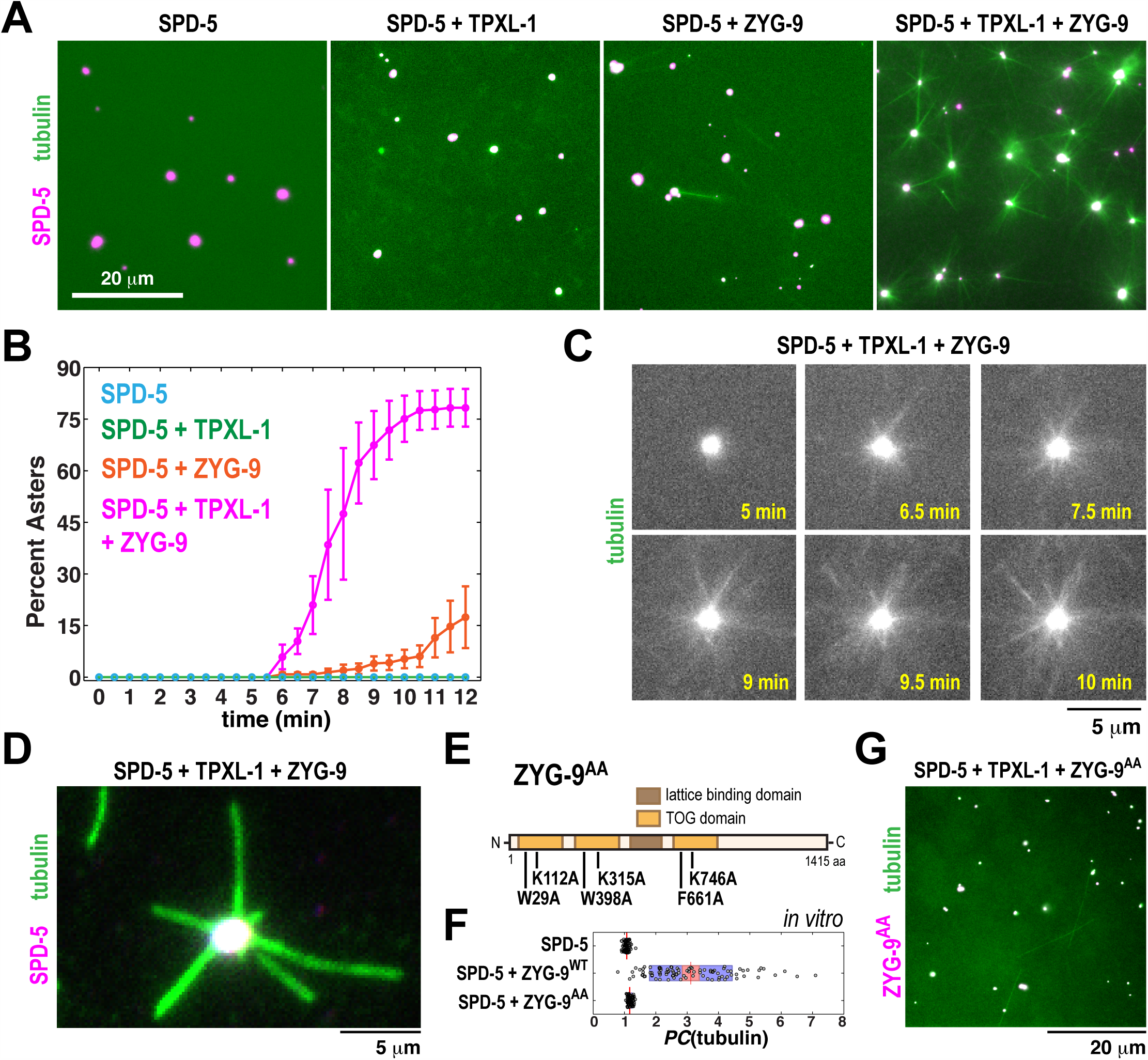
SPD-5 droplets containing TPXL-1 and ZYG-9 organize microtubule asters. A. 900 nM SPD-5 droplets (1:8 ratio of TagRFP-labeled to unlabeled) were incubated with 2.5 μM α/β tubulin (1:4 ratio of alexa488-labeled to unlabeled), 2 mM GTP, 100 nM TPXL-1, and 50 nM ZYG-9. Images were taken after 10 min incubation at 23°C.
B. Quantification of (A) showing the percentage of SPD-5 droplets that have produced asters in one field of view (n = 3 experiments per condition; data are mean ± SEM; > 50 droplets per experiment).
C. Time-lapse images of aster formation from a SPD-5/ZYG-9/TPXL-1 droplet. α/β tubulin signal is shown. See also Movie S1.
D. SPD-5/TPXL-1/ZYG-9 asters were diluted into 10 μM taxol + 7.5% PEG and imaged by total internal reflection microscopy.
E. Schematic of the ZYG-9^AA^ mutant.
F. 500 nM SPD-5::TagRFP droplets were incubated with 100 nM α/β tubulin (alexa488-labeled), 2 mM GTP, and 50 nM wild-type ZYG-9 (ZYG-9^WT^) or mutant ZYG-9 (ZYG-9^AA^). After 10 min, partition coefficients were measured for tubulin (n = 92 droplets).
G. Same as in (A), except with 50 nM ZYG-9^AA^. See also Figure S5.

## Discussion

In this study, we have shown that the coiled-coil protein SPD-5, which forms the PCM scaffold in *C. elegans*, assembles into micron-scale, spherical droplets in the presence of crowding agents *in vitro.* PCM client proteins, including microtubule-associated proteins, then partition into this droplet phase via interactions with SPD-5. While the SPD-5 scaffold becomes more stable over time, the clients remain loosely bound and mobile and are sufficient to recruit tubulin and form microtubule asters (Figure 7A,B). Thus, the PCM scaffold acts as a selective compartment into which clients partition. These conclusions are consistent with a theoretical framework describing PCM organization and growth (Zwicker et al., 2014).

### Pericentriolar material is a selective, viscous droplet

PCM must satisfy unique material and dynamic properties to function properly. First, PCM must be permeable: it must be porous enough to permit entry and diffusion of the correct proteins to allow biochemical reactions such as microtubule nucleation. Second, PCM must be selective: it must concentrate tubulin, microtubule-associated proteins, and regulatory proteins, and exclude noncentrosomal proteins. Finally, PCM must be both flexible and strong: it must be able to incorporate new material and expand in preparation for mitosis while resisting microtubule-dependent pulling forces. Internal rearrangement of PCM is also necessary to allow microtubules nucleated in the interior to polymerize and extend beyond the PCM. Our *in vitro* data suggest that the PCM satisfies these requirements by forming a viscous droplet by phase separation of the scaffold protein SPD-5 (Figure 7A).

**Figure 7.**
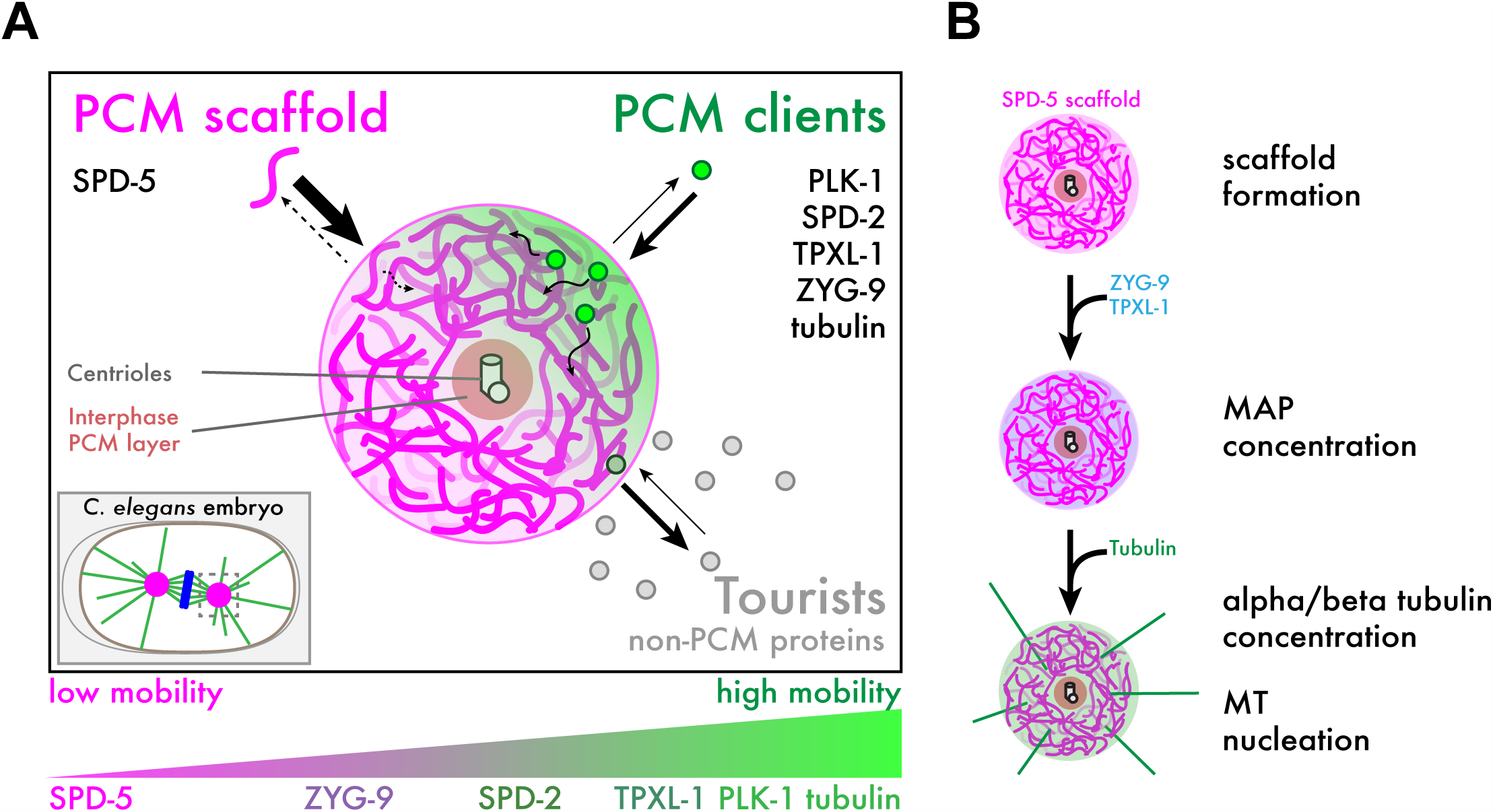
Proposed model for PCM organization and microtubule nucleation. A. Phase separation of SPD-5 around centrioles creates an amorphous, gel-like material (“PCM scaffold”, pink) that selectively concentrates proteins needed for centrosome function (“PCM clients”, green). Non-centrosomal proteins can enter and pass through the PCM scaffold but do not bind strongly (“Tourists”, gray). Arrow thickness indicates chemical equilibrium between the droplet phase and bulk solutions. Different PCM client proteins can exhibit different degrees of mobility within the PCM scaffold (bottom panel).
B. Proposed mechanism of PCM-based microtubule nucleation. The SPD-5 scaffold forms first, then concentrates microtubule-associated proteins (MAPs) such as TPXL-1 and ZYG-9. These MAPs concentrate tubulin within the scaffold phase and lower the energy barrier for tubulin nucleation to generate microtubule asters.

The idea that the PCM is a viscous droplet is supported by the following observations. First, both *in vitro* SPD-5 droplets and *in vivo* PCM are spherical. Second, *in vitro* SPD-5 droplets and *in vivo* PCM incorporate new material isotropically and expand isotropically (see also (Laos et al., 2015)). Third, PCM client proteins concentrate into *in vitro* SPD-5 droplets and *in vivo* PCM but rapidly rearrange within these compartments. Finally, *in vitro* SPD-5 droplets are amorphous and can even coalesce. With time, however, SPD-5 droplets no longer internally rearrange, exchange components between the droplet and bulk solution, nor coalesce *in vitro*, consistent with the dynamic properties of centrosomal SPD-5 in metaphase-arrested embryos (Laos et al., 2015). We do not yet know the material property of these hardened states. The decrease in dynamics could represent a transition into a jammed, glass-like state or solidification into a gel. Considering that PCM client proteins are still mobile within hardened SPD-5 droplets *in vitro* and mitotic PCM *in vivo*, it is likely that nanometer-wide pores exist, consistent with a gel-like state. This transformation could serve a functional purpose by first establishing PCM shape and allowing rapid PCM expansion early in the cell cycle, and then allowing the PCM to become rigid enough to resist MT-pulling forces during mitosis.

Many of the behaviors we describe for SPD-5 droplets have been seen in cellular bodies that form through liquid-liquid phase separation from cytoplasm, such as ribonucleoprotein granules, PML bodies, and FUS droplets (Banani et al., 2016; Brangwynne et al., 2009; 2011; Lin et al., 2015; Patel et al., 2015). Like SPD-5 droplets, these cellular bodies are spherical, exhibit liquid-like behaviors and even undergo liquid-to-solid conversion *in vitro*, albeit over vastly different time scales; for example, FUS droplets harden over the course of several hours, whereas SPD-5 hardens over several minutes. These phase-separated protein systems can also exhibit selective scaffolding properties as seen for SPD-5. Reconstituted nucleoporins form stable hydrogels that permit entry and exit of nuclear transport receptors while excluding other proteins (Frey and Görlich, 2007; Hülsmann et al., 2012; Schmidt and Görlich, 2015). Another example includes interactions between multivalent SUMO-SIM droplet scaffolds and their monovalent client proteins (Banani et al., 2016). In this system, the scaffold component is far more stable than the weakly bound client proteins, which rapidly exchange with the bulk solution. By analogy, we propose that macromolecular crowding and multivalent homotypic interactions drive phase separation of SPD-5 into a condensed, amorphous phase, which we call the “scaffold phase” (Figure 7A,B). SPD-2 and PLK-1 regulate PCM assembly by lowering the energy barrier for SPD-5 phase separation. Client proteins, such as TPXL-1 and ZYG-9, then partition into the SPD-5 phase to drive microtubule nucleation.

A unique feature of SPD-5 that sets it apart from other phase separating protein systems is domain architecture. Ribonucleoprotein granules, FUS, and nucleoporins mediate interactions through disordered, low-complexity domains and interactions with RNA molecules (Frey and Görlich, 2007; Kato et al., 2012; Patel et al., 2015). However, SPD-5 lacks long, highly disordered regions and canonical RNA binding domains and instead contains nine predicted coiled-coil domains which make up ~40% of the protein (Figure S6A). Coiled-coil domains have been well described in the formation of stoichiometric protein complexes. Our results raise the possibility that coiled-coil domains also mediate homotypic interactions to drive the formation of amorphous droplets. Interestingly, surface plasmon resonance experiments indicated that coiled-coil interactions occur in a biphasic manner, implying a fast, initial binding stage followed by a rate-limiting, rearrangement stage that can last >90s (De Crescenzo et al., 2003). This biphasic assembly of coiledcoils could represent a shift from weak to strong interactions and could potentially explain how SPD-5 droplets form as viscous liquids that quickly harden. It will be fascinating in the future to understand the relation between coiled-coil domains and SPD-5 phase separation.

### Macromolecular crowding in the cytoplasm could influence centrosome assembly

An unexpected finding from our work was that macromolecular crowding changed the organization of SPD-5 assemblies. Without crowding, SPD-5 assembles into dispersed networks that do not nucleate microtubules. Yet, when placed in a solution of PEG, Ficoll, Dextran, or Lysozyme, SPD-5 assembles into droplets that morphologically, functionally, and dynamically resemble *in vivo* PCM. These results raise the possibility that inert macromolecules in *C. elegans* embryo cytoplasm shape PCM into a spherical structure. This phenomenon is likely an entropic effect of crowding and not a chemical effect because: 1) different macromolecules (glycols, sugars, and protein) exert the same effect, and 2) these macromolecules must be in molar excess to exert this effect (e.g., mM concentration of PEG vs. nM concentration of SPD-5). Additionally, we found that SPD-5 droplet formation was sensitive to the size of the macromolecule; SPD-5 formed droplets in a solution of 9% PEG-3350 (MW: 3,350 Da) but not in 9% PEG-300 (MW: 300 Da). This result is consistent with a depletion attraction mechanism. Similar size-dependent crowding effects have also been shown to enhance actin polymerization (Drenckhahn 1986).

Depletion attraction is a well-described entropic effect that drives the coalescence of large molecules in a crowded solution of smaller molecules, called “depletants” (for a review, see (Marenduzzo et al., 2006)). As two large molecules approach each other, the space between them becomes inaccessible to the depletants, resulting in an unfavorable increase in entropy for the entire system. As a result, an inward osmotic force is generated that drives the two large molecules together and diminishes the excluded volume. For depletion attraction to occur, the smaller molecules must be in excess and in a certain size range relative to the larger, more rare molecules. The smaller the depletant is, the more likely it can access the space between the larger molecules, which eliminates the entropic penalty caused by exclusion. As the depletant approaches the size of the larger molecule, the amount of newly accessible space formed after the larger molecules coalesce becomes negligible and the depletion attractive force diminishes. In our system, PEG is in excess and could act as the depletant that drives the larger, limiting SPD-5 molecules together. Our results suggest that depletion attraction influences the nature of SPD-5’s homotypic interactions so that a droplet, rather than an extended network, is formed. It will important to determine how the more complex embryonic cytoplasm influences centrosome formation. So far, we have not been able to observe SPD-5 droplet formation in *C. elegans* extract due to the presence of an unidentified inhibitory agent (unpublished data).

### Tubulin concentration and microtubule stabilization drive aster formation

A key goal of this work was to identify the minimum factors required for PCM-dependent microtubule nucleation. We have shown that two proteins that affect microtubule dynamics, TPXL-1 (*C. elegans* homolog of TPX2) and ZYG-9 (*C. elegans* homolog of ch-TOG/XMAP215), can partition into SPD-5 droplets. These SPD-5/TPXL-1/ZYG-9 droplets concentrate tubulin ~4-fold and are sufficient to organize radial microtubule arrays. Therefore, an amorphous droplet of SPD-5, into which TPXL-1, ZYG-9 and tubulin partition, represents a minimal microtubule organizing center.

We do not yet know the detailed mechanism by which these two MAPs drive microtubule assembly. One possibility is that they simply concentrate tubulin into a dense phase, since microtubule nucleation occurs spontaneously at tubulin concentrations >20 μM *in vitro.* In our assays, we could achieve microtubule aster formation with as little as 2.5 μM tubulin, which translates to ~10 μM tubulin within SPD-5/TPXL-1/ZYG-9 droplets (Figure 5), a value that is too low to drive spontaneous microtubule nucleation. However, we cannot rule out the possibility that enriched tubulin subdomains exist within SPD-5 droplets, where tubulin concentrations could be above the critical limit for nucleation. On the other hand, Roostalu *et al.*, (2015) demonstrated that human TPX2 and chTOG synergistically enhance MT nucleation, thereby lowering the tubulin concentration needed for nucleation. Thus, we currently propose that PCM organizes microtubule arrays in part through complementary mechanisms of tubulin concentration and microtubule-stabilization by the TPXL-1/ZYG-9 module (Figure 7B).

RNAi and genetics experiments in *C.elegans* embryos have suggested that *in vivo*, PCM-based microtubule nucleation is very robust and involves multiple pathways. For example, double RNAi knockdown of ZYG-9 and TPXL-1 or single RNAi knockdown of gamma tubulin reduces, but does not fully eliminate, PCM-based microtubule nucleation *in vivo* (Figure S6B; (Hannak et al., 2002; Strome et al., 2001)). *In vitro* reconstitution experiments have shown that the TPX2/ch-TOG (TPXL-1/ZYG-9) module and gamma tubulin complexes each promote microtubule nucleation, likely through stabilization of spontaneously-formed nuclei versus templating new growth (Kollman et al., 2015; Roostalu et al., 2015; Wieczorek et al., 2015). These findings suggest that robustness is guaranteed through several overlapping, but mechanistically distinct microtubule nucleation pathways. There may even be additional, unexplored pathways; PCM-based microtubule nucleation still occurred even after triple inhibition of ZYG-9, TPXL-1, and gamma tubulin *(zyg-9^ts^;tpxl-1(RNAi);tbg-1(RNAi);* unpublished data). Future work *in vivo* and *in vitro* is required to determine the relative contributions of these and additional microtubule-associated proteins to aster formation.

### Insights into PCM nucleation

In mitotically dividing cells, PCM forms only around centrioles to ensure bipolar spindle formation. How is the location of PCM assembly regulated? Previous work that employed fusion experiments between dividing and non-dividing cells in the *C. elegans* gut suggested that activated SPD-2 initiates PCM formation (Yang and Feldman, 2015). During fertilization, SPD-2 initially localizes to the centrioles provided by the sperm, again suggesting that SPD-2 acts as the trigger for PCM formation (Kemp et al., 2004; Pelletier et al., 2004). Our *in vitro* experiments support this idea by showing that dense SPD-2 scaffolds nucleate SPD-5 droplets. Based on these studies and our results, it seems likely that activation of the SPD-2 layer coating interphase centrioles is the key step in initiating mitotic PCM formation. The cytoplasm, then, must provide crowding conditions that are unsuitable for spontaneous PCM formation. PEG concentrations > 4% drive SPD-5 on its own into droplets *in vitro.* However, a nucleation module (SPD-2/PLK-1) greatly lowers this requirement for PEG to 1.5%. Thus, we propose that embryonic cytoplasm more closely resembles the low-PEG state *in vitro* where droplet formation is regulated. It is likely that most phase separation systems are regulated through nucleation *in vivo*, which could explain why these systems sometimes require large amounts of crowding agents to form droplets *in vitro.*

### Comparison to other species

Many of the proteins that regulate PCM assembly and microtubule nucleation are conserved across diverse eukaryotic species, suggesting that the mechanisms of assembly and aster formation could be conserved as well. Homologues of PLK-1 and SPD-2 are required for PCM assembly in *Drosophila* and human cells (Conduit et al., 2014a; Giansanti et al., 2008; Haren et al., 2009; Lee and Rhee, 2011; Sunkel and Glover, 1988; Zhu et al., 2008), and homologues of TPXL-1 and ZYG-9 are required for full microtubule nucleation from centrosomes in numerous metazoans (Barr and Bakal, 2015; Gergely et al., 2003; Goshima, 2011; Gruss et al., 2001; Popov et al., 2002; Wittmann et al., 2000). While there are no sequence-based homologues of SPD-5 outside of nematodes, there is mounting evidence that functional homologues exist, such as Centrosomin/CDK5Rap2 and DPLP/ Pericentrin. In human cells, PCM assembles through Polo kinase-regulated accumulation of Pericentrin around centrioles (Lee and Rhee, 2011). The same has been shown for Centrosomin in *Drosophila* (Conduit et al., 2010). Furthermore, *in vitro* analysis of a Centrosomin fragment indicated that phosphorylation promotes homo-oligomerization (Conduit et al., 2014a). Thus, Polo kinase-regulated formation of a scaffold that recruits PCM clients is a common mechanism for PCM assembly.

Several lines of experimentation also indicate that PCM assembly may occur through phase separation in other organisms. The most compelling evidence comes from a comprehensive study of protein-protein interactions within the *Drosophila* centrosome using yeast-two-hybrid analysis (Galletta et al., 2016). This study revealed that PCM proteins make numerous heterotypic and homotypic interactions; for example, one Centrosomin fragment participated in 24 unique interactions. This high degree of multivalency should be sufficient to drive phase separation (reviewed in (Bergeron-Sandoval et al., 2016)). Additionally, the conserved spherical morphology of the centrosome is suggestive of phase separation. It is possible that centrosome shape is determined by a biophysical parameter such as surface tension. The next step will require a combination of *in vivo* dynamics studies and *in vitro* reconstitution to determine if centrosomes in other species display liquid-like properties as seen for *C. elegans* centrosomes.

## Author Contributions

J.B.W. and A.A.H. conceived the project. J.B.W. wrote the manuscript and performed all of the experiments, except those specifically attributed to other authors. B.F.G. performed some of the *in vitro* FRAP experiments and droplet assays. P.O.W. helped with protein purification and with initial characterization of the aster assays. J.M. performed cryo EM.

## Acknowledgements

We thank the Protein Expression & Purification and Light Microscopy facilities of MPI-CBG (Dresden). We thank Raul Grover for help with TIRF imaging; Andrea Zinke and Anne Schwager for help with worm maintenance; and Andrei Pozniakovsky for help with cloning DNA constructs. This project was funded by the Max Planck Society and the European Commission's 7th Framework Programme grant (FP7-HEALTH-2009-241548/MitoSys) and a MaxSynBio grant to A.H. J.B.W. was supported by an EMBO fellowship and MaxSynBio. J.M. was supported by EMBO and HFSP postdoctoral fellowships.

## SUPPLEMENTAL MATERIAL

### MATERIALS AND METHODS

#### Protein expression and purification

All expression plasmids are listed in Table S1. SPD-5, SPD-2, and PLK-1 proteins were expressed and purified as previously described (Woodruff and Hyman, 2015; Woodruff et al., 2015), with the following exception: SPD-2 was stored in its uncleaved form (MBP-TEV-SPD-2) and then thawed and treated with TEV protease for 1 hr prior to daily use. Porcine tubulin was used throughout this study and was purified as previously described (Gell et al., 2011). A coomassie-stained gel of proteins used in this study is shown in Figure S1D.

Full-length *zyg-9* (4245 bp), and *tpxl-1* (1521 bp), genes lacking stop codons were amplified from *C. elegans* cDNA by PCR and inserted into in-house-designed baculoviral expression plasmids (pOCC series). These proteins were expressed in SF+ insect cells and harvested 72 hr post infection. Cells were collected, washed, and resuspended in harvest buffer (50 mM Tris HCl, pH 7.4, 150 mM NaCl, 30 mM imidazole, 1% glycerol) + protease inhibitors (1 mM PMSF, 100 mM AEBSF, 0.08 mM Aprotinin, 5 mM Bestatin, 1.5 mM E-64, 2 mM Leupeptin, 1 mM Pepstatin A)(Calbiochem) and frozen in liquid nitrogen. All subsequent steps were performed at 4°C. Cells were lysed using a dounce homogenizer. CHAPS detergent was added to a final concentration of 0.1%, and NaCl was added to a final concentration of 500 mM. The crude lysate was clarified by centrifugation for 25 min at 146,000 × g.

EB1-GFP was expressed in *Escherichia coli* (BL21(DE3) CodonPlus-RIL) induced with 0.5 mM IPTG for 16 h at 18°C. Cell pellets were frozen in liquid nitrogen the resuspended in EB1 lysis buffer (50 nM HEPES, pH 7.2, 400 mM NaCl, 2 mM MgCl_2_, 20 mM imidazole, 1 mM DTT, 0.1% CHAPS) + protease inhibitors. Cell were lysed by two passages through an Avestin Emulsiflex-C5 then clarified by centrifugation for 25 min at 146,000 × g.

To purify constructs, the clarified lysate was incubated with Ni-NTA agarose (Qiagen) for 2 hr. The agarose beads were washed with 10 column volumes of wash buffer (50 mM Tris HCl, pH 7.4, 500 mM NaCl, 30 mM imidazole, 1% glycerol, 0.1% CHAPS), and the protein was eluted with 250 mM imidazole. Proteins were incubated with PreScission protease (ZYG-9 and TPXL-1 constructs) or TEV protease (EB1-GFP) overnight to remove affinity tags. The proteins were then purified over a Superdex 200 HR 10/30 gel filtration column (GE Healthcare Life Sciences) equilibrated with storage buffer (50 mM Tris HCl, pH 7.4, 500 mM NaCl, 0.5 mM DTT, 1% glycerol, 0.1% CHAPS) using an AKTA Pure FPLC system (GE Healthcare). Peak fractions were pooled and diluted into low salt (final salt concentration = 50 mM NaCl), then loaded onto a SP Sepharose column. The column was washed stepwise with increasing salt concentrations and eluted with storage buffer containing 10% glycerol. Proteins were then aliquoted in PCR tubes, flash-frozen in liquid nitrogen, and stored at −80°C. Protein concentration was determined by measuring absorbance at 280 nm using a NanoDrop ND-1000 spectrophotometer (Thermo Scientific).

#### *In vitro* SPD-5 droplet assembly, imaging, and analysis

SPD-5 droplets were formed by adding concentrated SPD-5::GFP or SPD-5::TagRFP to Droplet buffer (25 mM HEPES, pH 7.4, 150 mM KCl) containing polyethylene glycol (SIGMA) and fresh 0.5 mM DTT. For most experiments, SPD-5 droplets were visualized with an inverted Olympus IX71 microscope using 60x 1.42 NA or 100x 1.4 NA Plan Apochromat oil objectives, CoolSNAP HQ camera (Photometrics), and DeltaVision control unit (AppliedPrecision). Images were also taken using an inverted Olympus IX81 microscope with a Yokogawa spinning-disk confocal head (CSU-X1), a 100x 1.4 NA Plan Apochromat oil objective, and an iXon EM + DU-897 BV back illuminated EMCCD (Andor).

Images were analyzed in MATLAB or FIJI. In brief, particles were identified through applying a threshold then using the particle analyzer function in FIJI. When analyzing droplet formation, we report the sum of the integrated intensities of each droplet per image (total droplet mass). When analyzing client protein recruitment to SPD-5 droplets, we report the ratio of the mean intensity of the client protein in the droplet vs. the bulk solution after background subtraction for individual droplets (partition coefficient, or *PC*).

#### High-throughput automated droplet imaging and analysis

Two separate solutions containing 50-200 nM SPD-5 in Droplet buffer and 6-25% PEG-3350 in Droplet buffer were dispensed into two rows of a 96 well plate. A pipetting robot (Tecan, Model Freedom EVO 200) equipped with a TeMo 96 pipetting head with disposable tips (50 ul) was used to mix the SPD-5 and PEG solutions 1:1 and then transfer each condition in quadruplicate to a 384 well plastic bottom imaging plate (15 μl per well) to synchronize droplet formation. The 384 well plate was centrifuged for 5 min at 4,000 rpm to sediment droplets, then imaged using a Yokogawa CV7000 high-content spinning disk confocal microscope equipped with a 60x 1.2 NA water immersion objective. Z-stacks (3 planes, 0.2 μm spacing) were taken at 6 different positions per well, then compressed using a maximum intensity projection (3 × 6 × 4 = 72 images total per condition; 72 × 60 = 4320 images total per plate). Images were segmented with CellProfiler 2.1 and analyzed using KNIME 2.11. The heat maps in Figures 1B and S1C were generated with the KNIME plugin ‘HCS Tools’ with R script and represents the mean values of the 4 wells assigned to the designated condition.

#### Fluorescence Recovery After Photobleaching (FRAP)

For FRAP analysis of *in vitro* SPD-5 droplets, samples were mounted in an imaging chamber made with a PEG-functionalized coverslip, double-sided tape, and a microscope slide. To prepare the coverslips, coverslips were incubated in 100% ethanol for 20 min in a sonicating water bath, then washed with MilliQ H_2_O. The coverslips were then incubated overnight in a solution containing 250 mL Toluene, 575 μL 3-[Methoxy(polyethyleneoxy)6-9propyl]trimethoxysilane, tech-90 (abcr GmbH), and 200 μL HCL (37%). Finally, the coverslips were washed twice with 100% ethanol and MilliQ water and dried with compressed air. Coverslips were stored in a desiccation chamber.

SPD-5 droplets were formed in buffer containing 9% PEG, 100 nM NaCl, 345 mM KCl, 0.02% CHAPS, and oxygen scavengers (40 mM Glucose (Sigma), 130 μg/ml Glucose oxidase (Sigma), and 24 μg/ml Catalase (Sigma). After 30s, these droplets were diluted 1:1 into a solution containing the relevant GFP-tagged client protein diluted in water. Droplets were then loaded into the imaging chamber after 10 min of incubation at 23°C.

FRAP experiments were performed on the same spinning disk confocal microscope mentioned above using a 100x 1.4 NA oil immersion objective. Laser micro-irradiation were performed with a 488 nm or 405 nm laser (5.5 mW-11.0 mW output) with a 50 ms exposure time and 1 × 1 binning. Droplets were bleached for 15 ms and images were taken at 1 s intervals. Analysis of the recovery curves and the half-time recovery were carried out with the FIJI/ImageJ macro (http://imagej.net/Analyze FRAP movies with a Jython script) and MATLAB.

FRAP measurements of *in vivo* centrosomes were performed under similar conditions with the following exceptions. Embryos were mounted on agar pads in M9 buffer and imaged using 488 nm or 561 nm lasers and a 60x 1.2 NA water immersion objective. Z-stacks (16 images spanning 33 μm) were collected every 5 sec with 2 × 2 binning. Maximum intensity projections of the Z-stacks were analyzed as above. Partial bleaching of PLK-1::GFP-labeled centrosomes could not be performed due to insufficient signal intensity and area. For nocodazole treatment of embryos, L4 worms were grown on *perm-1(RNAi)* feeding plates at 20°C for 16-18 hr, then dissected in an open imaging chamber filled with osmotic support medium (Carvalho et al., 2011; Wueseke et al., 2016) and 20 μg/ml nocodazole (Sigma).

#### Microtubule aster formation and imaging

To form asters, 900 nM SPD-5 droplets were assembled in Droplet buffer including 7.5 % PEG, 0.5 mM DTT, 100 nM TPXL-1, 50 nM ZYG-9, 2 mM GTP and a 2.5 μM (final) mix of Alexa488-labeled and unlabeled tubulin (1:4 molar ratio). Droplets were squashed under pre-cleaned cover slips and imaged as described above.

For TIRF imaging, droplets were assembled in the same way, except that Cy5-labeled and unlabeled tubulin (1:3 molar ratio) were used. After 10 min of incubation, asters were diluted into Droplet buffer containing 7.5% PEG, 10 mM taxol, 40 mM Glucose (Sigma), 130 μg/ml Glucose oxidase (Sigma), and 24 μg/ml Catalase (Sigma), then loaded into a pre-blocked flow chamber. Images were obtained using a Nikon Eclipse Ti microscope equipped with Perfect Focus System, a 1.49 NA PlanApo 100X oil immersion objective, a monolithic laser combiner (Agilent MLC 400), and an electron multiplying charge-couple device (EMCCD) camera (iXon ultra EMCCD, DU-897U, Andor). Illumination settings were as follows: 647-nm laser line at 5 mW, with a Cy5 filter set (exec: 642/20. Dichroic LP 647, em: 700/75) with 100 ms exposure time.

#### Cryo EM

4 μl of 1 μM untagged SPD5 in 9% PEG was deposited on glow discharge Cupper Quantifoil grids (R2/1, Cu 200 mesh grid, Quantifoil Micro Tools) and plunge-frozen into liquid ethane/propane mixture at close to liquid nitrogen temperature using a Vitrobot^®^ Mark 4 (FEI). The blotting conditions were set to blot force 0, 3.5 s blot time and 2 s drain time. Cryo-transmission electron microscopy observations were performed on a Titan Krios operated at 300 kV (FEI) equipped with a field-emission gun, a Quantum post-column energy filter (Gatan), and a special heated phase plate holder (FEI). Data was recorded on a K2 Summit (Gatan) direct detector camera operated in dose fractionation mode. Images were collected using SerialEM at 42000x EFTEM magnification, corresponding to a pixel size 0.421 nm, a Volta phase plate at zero target defocus and 10 e^−^/Å^2^.

#### Worm strains and RNAi

*C. elegans* worm strains were maintained following standard protocols. Worm strains used in this study are listed in Supplemental Table S2.

**Supplemental Table S1.**
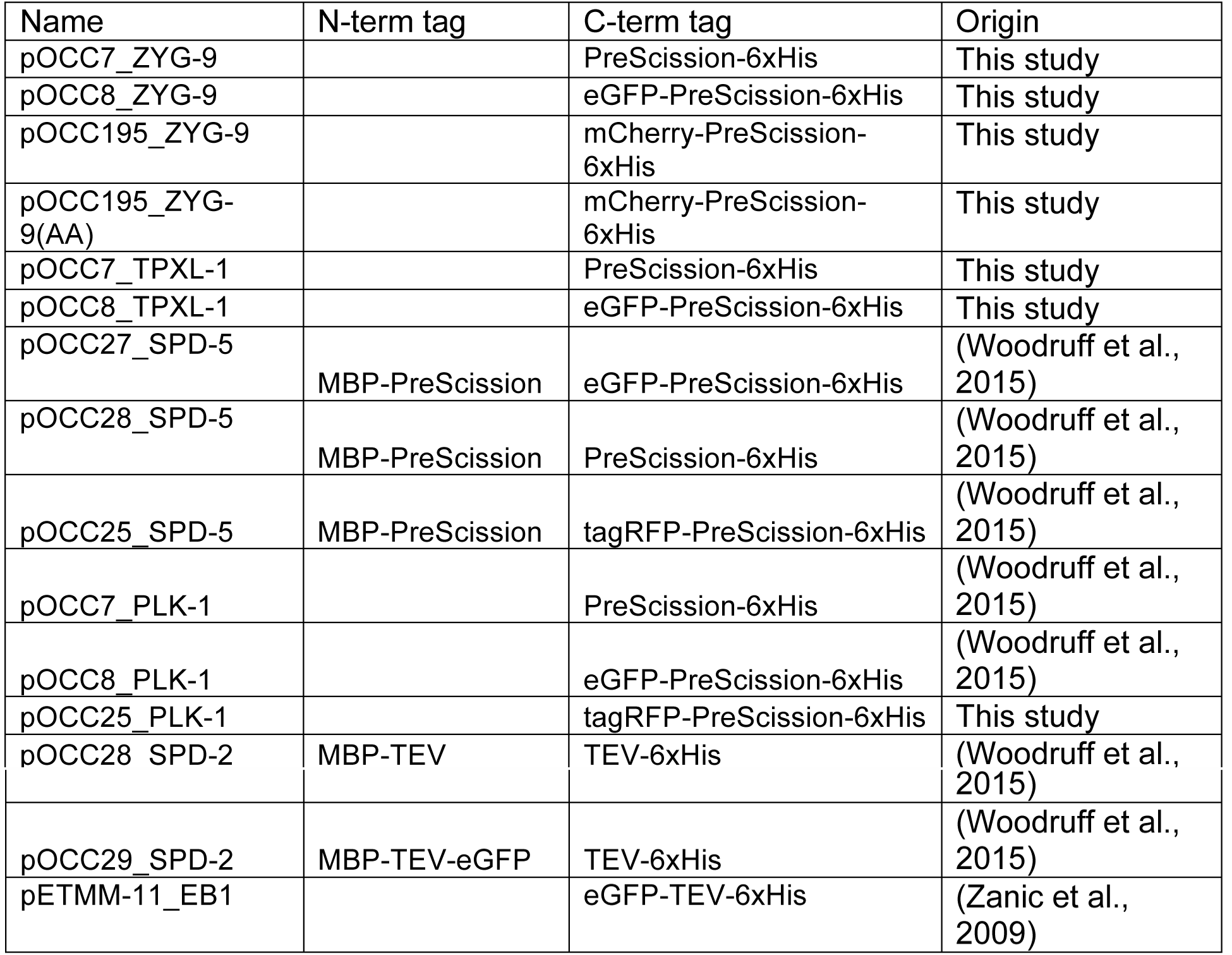
Protein expression plasmids.

**Supplemental Table S2.** *C. elegans* strains used in this study.

| Strain name | Main Marker | genotype |
| --- | --- | --- |
| OD847 | GFP::SPD-5 | unc-119(ed9) III; ltSi202[pVV103/ pOD1021; Pspd-2::GFP::SPD-5 RNAiresistant; cb-unc-119(+)]II |
| TH360 | PLK-1::GFP | unc-119(ed3)III; ddIs198[pie-1p::PLK-1b(synthetic introns, CAI 0.7)::GFP; unc-119(+)] |
| TH53 | TPXL-1::GFP | unc-119(ed3)III; ddIs12[pie-1p::TPXL-1::GFP;unc-119(+)] |
| TH165 | ZYG-9::mCherry | unc-119(ed3)III; ddIs52[WRM0614bG11 GLCherry::zyg-9;Cbr-unc-119(+)] |
| TH355 | SPD-2::GFP | unc-119(ed3)III; ddIs195[pie-1p::SPD-2(genomic introns, CAI 0.37)::GFP; unc-119(+)] |
| TH154 | GFP::TBB-2 | unc-119(ed3) ruls57[pAZ147: pie-1p/GFP::C36E8.5] III |

**Figure S1.**
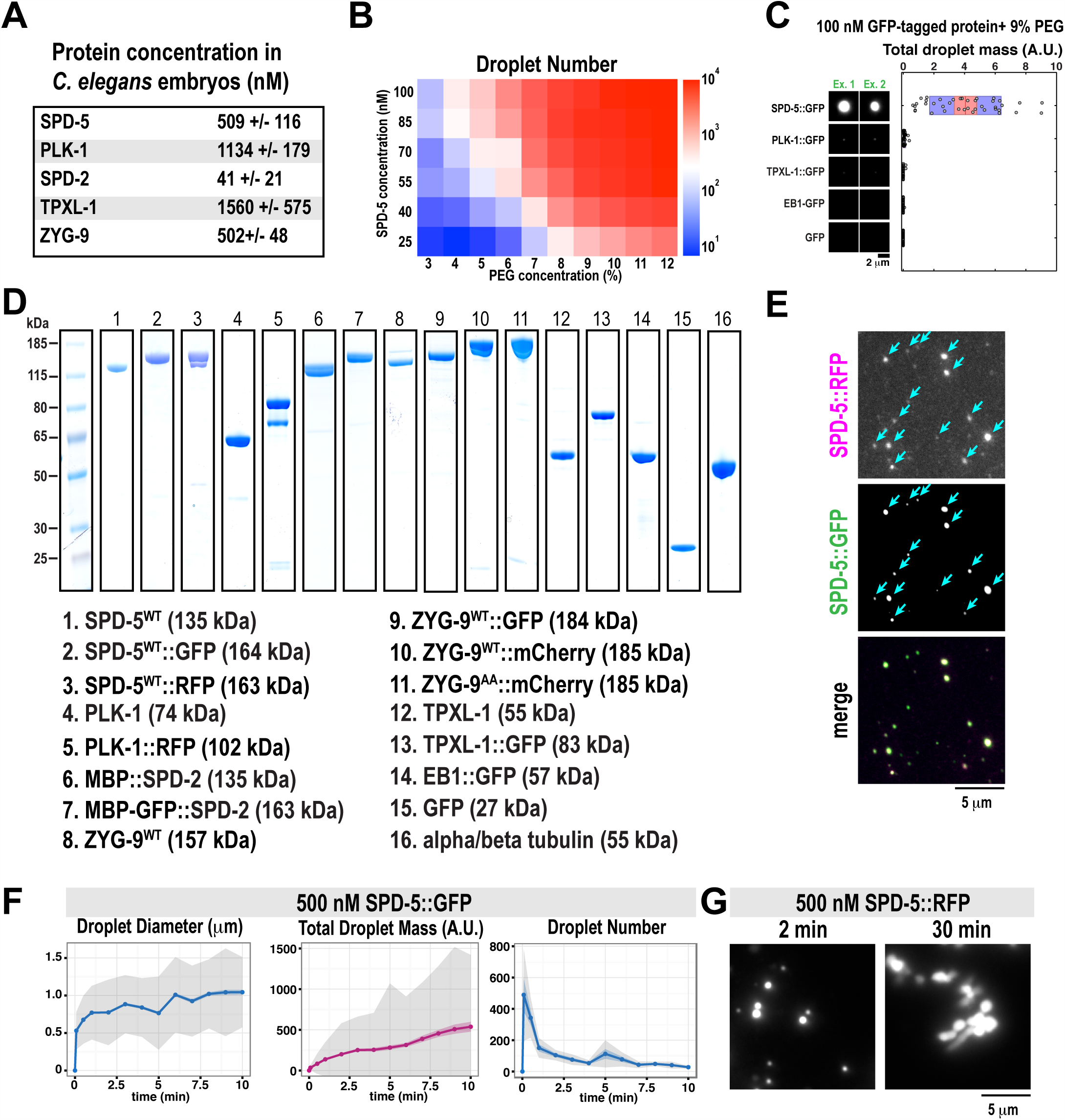
Analysis of PCM proteins in different conditions *in vitro*. A. Protein concentrations of indicated proteins in *C. elegans* embryos (mean ± S.D.; n = 3; see (Saha et al., 2016)).
B. Average number of droplets identified per condition from the experiment in Figure 1C.
C. Different GFP-tagged proteins (100 nM) were incubated for 5 min in 9% PEG. Only SPD-5::GFP formed detectable droplets. Total droplet mass was measured per field of view (n= 40 fields of view per condition). Two representative images are shown per condition.
D. Coomassie-stained gels showing the proteins used in this study.
E. Larger field of view from the seed expansion experiment in Figure 1D. SPD-5::GFP incorporates into existing SPD-5::TagRFP seeds rather than forming new droplets.
F. 500 nM SPD-5::GFP was incubated in 9% PEG and analyzed over time. Droplet diameter, total mass per image, and droplet number are shown (data are mean (lines), 95% confidence intervals (colored shaded area), and S.D. (grey shaded area); n = 11 images per time point).
G. SPD-5 droplets stop fusing over time and tend to stick together, forming large, amorphous structures (see 30 min).

**Figure S2.**
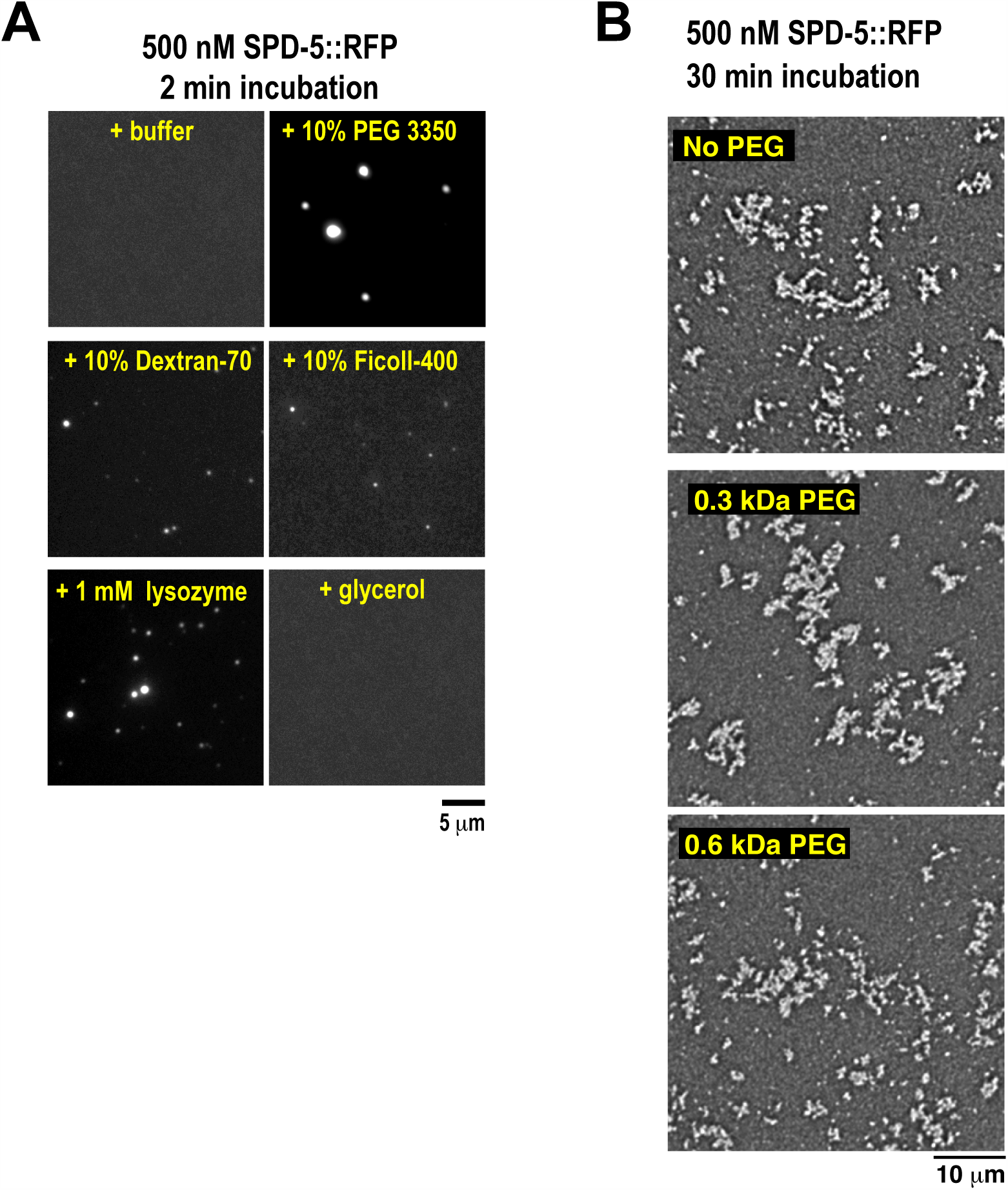
Analysis of SPD-5 assembly in different crowding conditions *in vitro*. A. 500 nM SPD-5::TagRFP was incubated for 2 min with indicated crowding agents.
B. Reactions were prepared as in Figure 2. 500 nM SPD-5::TagRFP forms networks after 30 min in conditions not conducive to droplet formation.

**Figure S3.**
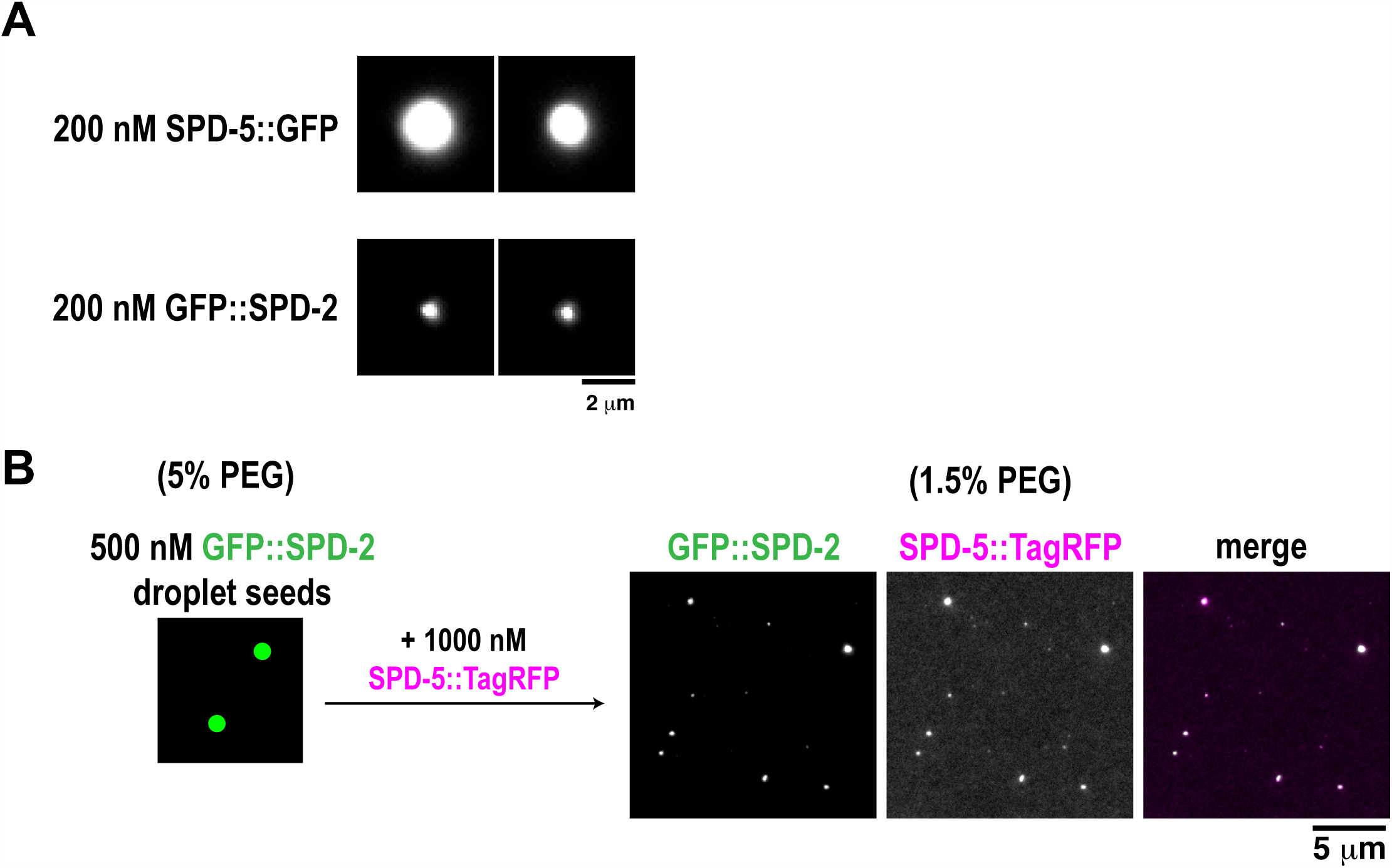
SPD-2 seeds are supramolecular scaffolds that nucleate SPD-5 droplets. A. Comparison of 200 nM SPD-5::GFP with 200 nM GFP::SPD-2 in a 5% PEG solution after a 10 min incubation at 23°C. Two representative droplets are shown.
B. 500 nM GFP::SPD-2 was incubated in 5% PEG for 5 min to form droplet seeds. These seeds were diluted into a solution containing no PEG and 1000 nM SPD-5::TagRFP so that the final concentration of PEG was 1.5%.

**Figure S4.**
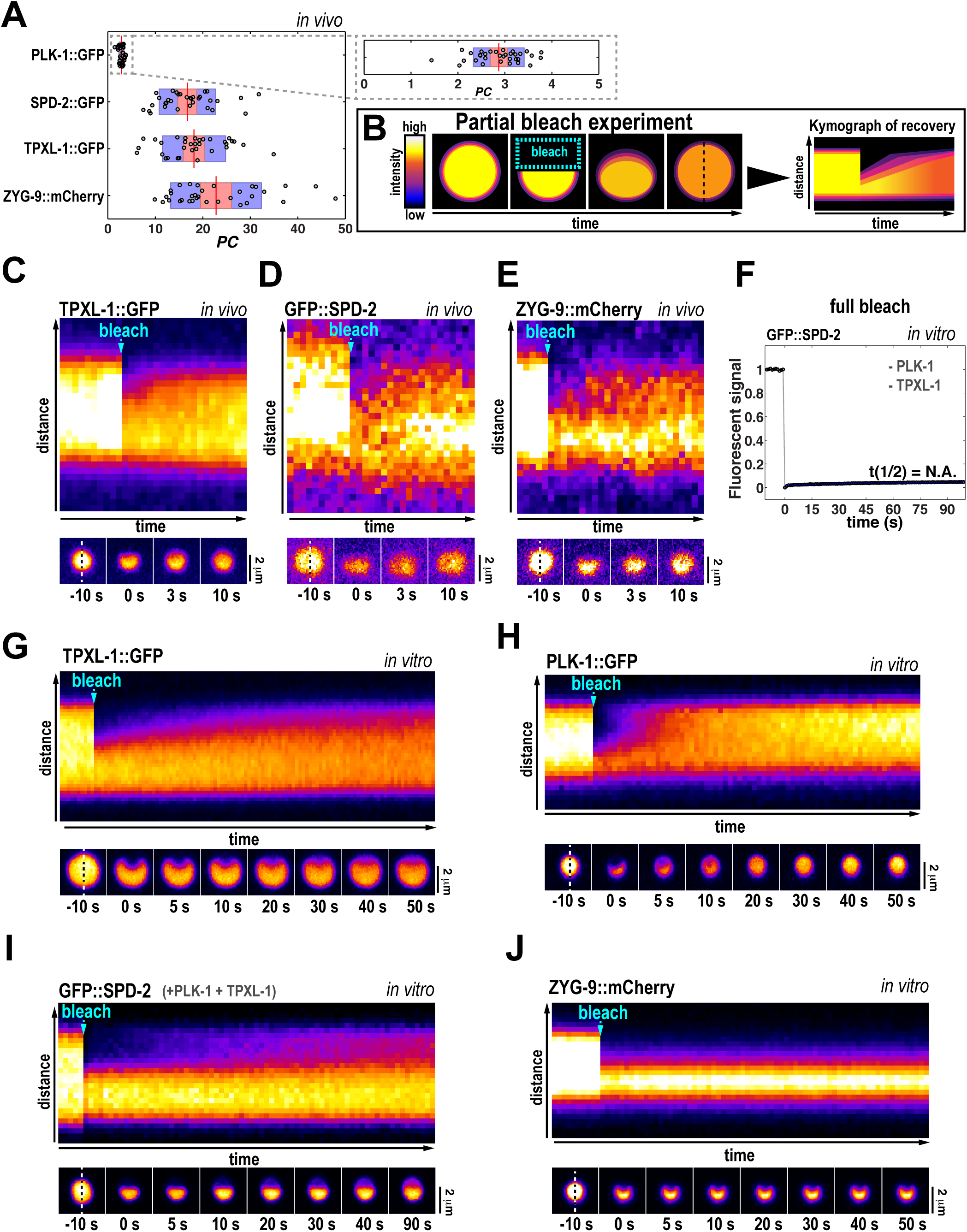
Analysis of PCM client protein recruitment and mobility. (A) Partition coefficients for PCM client proteins in 1-4 cell stage embryos during mitosis. Red bars represent the mean, red shaded areas represent 95% confidence intervals, and blue shaded areas represent one standard deviation (n = 33 centrosomes per strain). The gray box is a magnification of the PLK-1::GFP data.
(B) Schematic of the partial bleach experiments. Images are pseudocolored (high intensity to low intensity: white, yellow, red, purple, blue, black) to improve contrast. For analysis, a kymograph is made along the dotted line.
(C-E) Partial bleaching of TPXL-1::GFP (C), GFP::SPD-2 (D), and ZYG-9::mCherry (E) signal *in vivo.* Kymograph is along the dotted line.
(F) Analysis of GFP::SPD-2 signal in SPD-5 droplets after the entire droplet was bleached (same as in Figure 4E, except without unlabeled PLK-1 and TPXL-1)(n=15).
(G-J) Partial bleaching of TPXL-1::GFP (G), PLK-1::GFP (H), GFP::SPD-2 (I), and ZYG-9::GFP (J) signal in SPD-5 droplets *in vitro*. Kymograph is along the dotted line.

**Figure S5.**
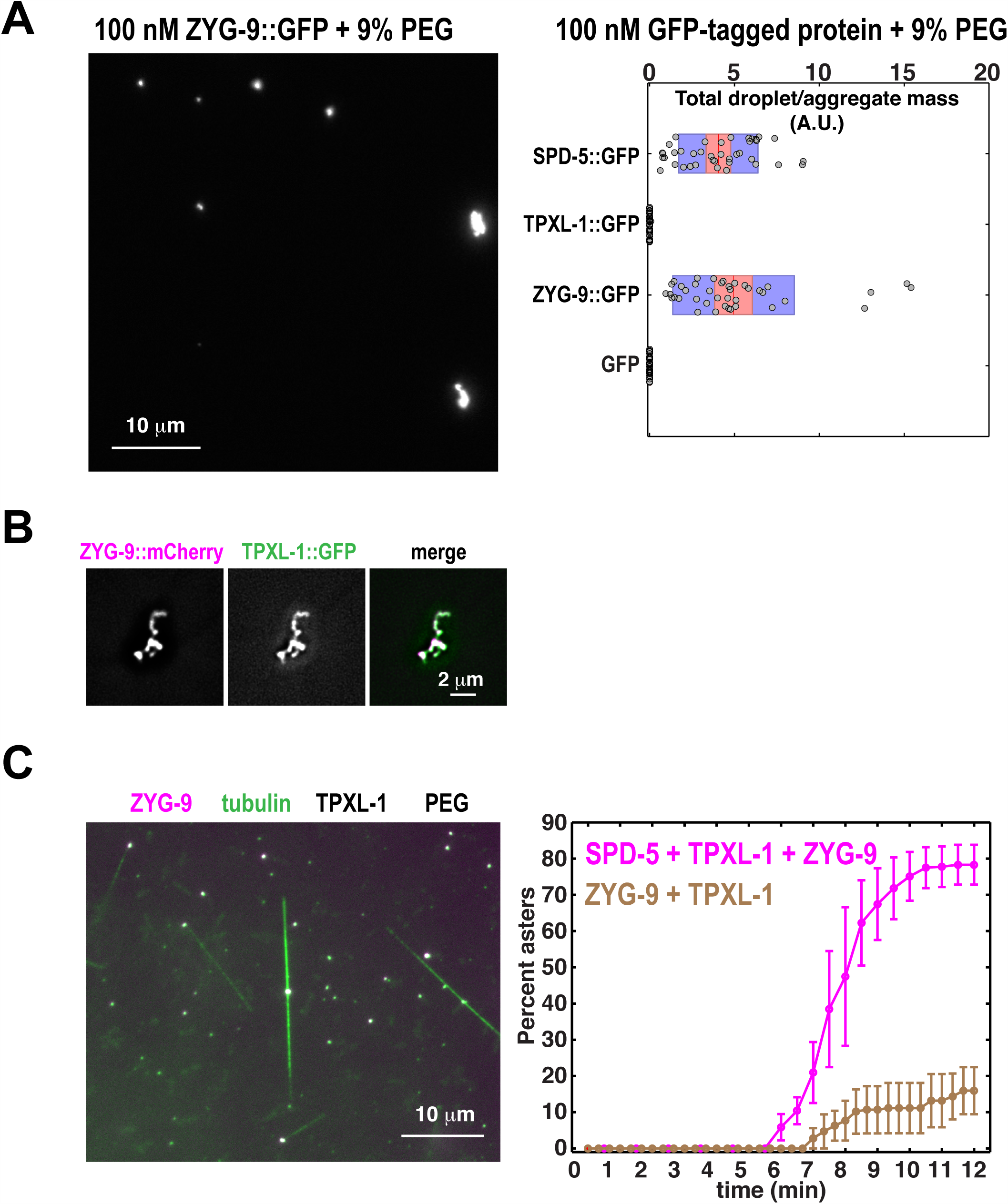
ZYG-9 aggregates are weak scaffolds that are insufficient to organize robust microtubule asters. A. 100 nM SPD-5::GFP, TPXL-1::GFP, ZYG-9::GFP, and GFP were incubated for 10 min at 23°C in a 9% PEG solution, then analyzed as in Figure S1C. ZYG-9 forms aggregates rather than droplets (n = 40 images analyzed per condition).
B. 500 nM ZYG-9::mCherry was incubated for 1 min at 23°C in a 9% PEG solution, then 100 nM TPXL-1::GFP was added. Images were taken 9 min later.
C. Same experiment as in Figure 6A, except SPD-5 was excluded from the reactions. The graph depicts the percentage of ZYG-9 aggregates (brown line) or SPD-5 droplets (magenta line) that have nucleated microtubules in one field of view (n = 3 experiments per condition; > 50 droplets per experiment; mean ± SEM).

**Figure S6.**
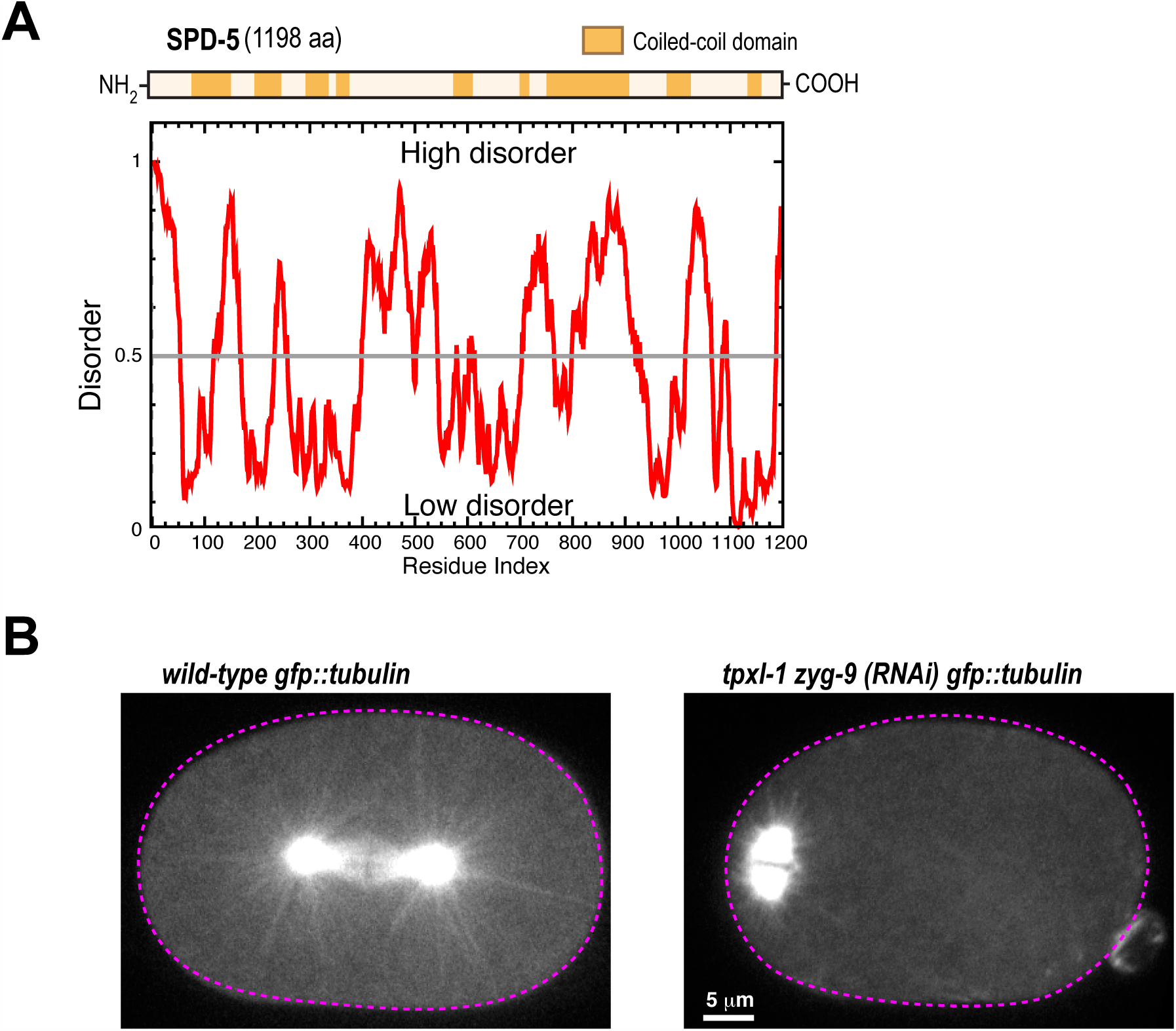
SPD-5 properties and *in vivo* analysis of microtubule nucleation. A. Domain organization of SPD-5 showing coiled-coil domains (top, analysis with MARCOIL at 90% threshold (Alva et al., 2016)) and disordered regions (bottom, analysis with PONDR-FIT (Xue et al., 2010)).
B. Mitotic *C. elegans* embryos expressing GFP::tubulin. Double *tpxl-1 zyg-9 (RNAi)* reduces spindle length and number of centrosome-based microtubules. Cell outline is in magenta.

